# Promoter-associated RNA polymerase III shapes RNA polymerase II-dependent inflammatory gene expression during viral infection

**DOI:** 10.64898/2026.07.13.738346

**Authors:** Azra Lari, Sahil B. Shah, Sarah Batarseh, Mikkel Nagorsen, Britt A. Glaunsinger

## Abstract

Cells must be primed to rapidly induce inflammatory gene expression upon infection while also tuning the level of induction to avoid immunopathology. Here, we identify RNA polymerase III (Pol III), best known for transcribing noncoding RNAs, as a dual-function regulator of RNA polymerase II (Pol II)-dependent inflammatory gene expression. Pol III is selectively enriched at promoters of innate immune, pro-inflammatory, and stress-response genes, where it maintains chromatin accessibility and supports basal transcription. Upon infection with murine gammaherpesvirus 68 (MHV68), Pol III redistributes from these promoters to retrotransposon loci, coinciding with enhanced expression of inflammatory genes. Depletion of the Pol III transcription factor Brf1 further amplifies inflammatory transcription during infection with MHV68, herpes simplex virus-1, and influenza A virus. Genes restrained by Pol III have TATA-box-enriched promoters and are functionally dependent on TATA-binding protein (TBP), suggesting that Pol III modulates inflammatory gene expression by competing with Pol II for shared transcriptional machinery. Thus, Pol III is a chromatin licensor in uninfected cells and an inflammation rheostat during viral infection.

**GRAPHICAL ABSTRACT:** 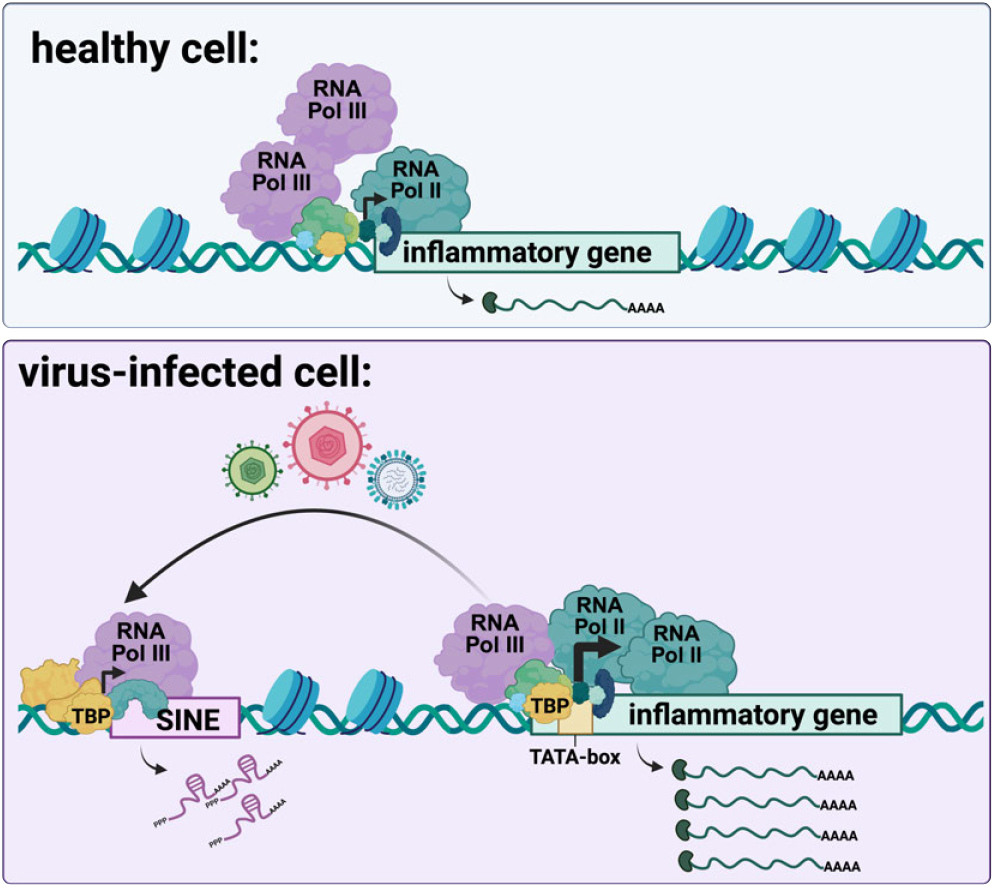

## INTRODUCTION

Inflammation associated with cellular antiviral defenses must be both rapid and restrained. Cells detect viral infection through pattern-recognition receptors, triggering rapid, tightly coordinated transcriptional programs that activate interferon-stimulated genes, pro-inflammatory cytokines, and innate immune effectors (Bartok and Hartmann, 2020; Li and Wu, 2021). These responses are coordinated through multiple regulatory layers, including both transcriptional and post-transcriptional control of gene expression (Medzhitov and Horng, 2009; Smale, 2010). Tempering pro-inflammatory pathways in response to stress and infection is also crucial, as unresolved inflammation contributes to pathogenic outcomes, including chronic inflammatory diseases associated with viral infections (Liu et al., 2017).

RNA polymerase II (Pol II) transcribes protein-coding genes, including those that encode pathogen-inducible innate immune and pro-inflammatory factors. However, evidence suggests that RNA polymerase III (Pol III) also plays a role in shaping the host’s innate immune response in mammalian cells (Graczyk et al., 2015; Jarrous and Rouvinski, 2021; Naesens et al., 2023; O’Neill, 2009). Pol III has been shown to transcribe sequences of viral origin, particularly those of common DNA viruses such as cytomegalovirus, vaccinia, herpes simplex virus-1, Epstein-Barr, and varicella zoster virus, generating immunostimulatory RNA ligands that can be detected by the cytoplasmic RNA sensor, RIG-I (Ablasser et al., 2009; Chiu et al., 2009; Ogunjimi et al., 2017). Mutations in Pol III subunits also underlie severe varicella-zoster virus infections (Carter-Timofte et al., 2018; Ogunjimi et al., 2017).

More canonically, Pol III transcribes a variety of cellular housekeeping noncoding RNAs (ncRNAs) essential for cellular function, including tRNAs, 5S rRNA, and U6 RNA. It also transcribes a class of ubiquitous noncoding retrotransposon genes in mammalian genomes known as short interspersed nuclear elements (SINEs). While typically silenced in healthy somatic cells to prevent insertional mutagenesis, SINE transcription is rapidly and robustly activated by Pol III in response to viral infection, cellular stress, inflammation, and cancer (Di Ruocco et al., 2018; Naesens et al., 2023; Willis and Moir, 2018). SINE-derived ncRNAs can stimulate innate immune signaling, largely through structural features such as extensive secondary structure and double-stranded RNA regions that enable recognition by innate immune sensors (Ahmad et al., 2018; Aune et al., 2022; Liddicoat et al., 2015). Other infection-inducible ncRNAs transcribed by Pol III, including pre-tRNAs and tRNA fragments, have also been shown to functionally impact gene expression in various cellular contexts (Dremel et al., 2023; Manning et al., 2025; Tucker et al., 2020).

A growing body of work has also established crosstalk between Pol II and Pol III as an important and previously underappreciated layer of transcriptional regulation, mediated through shared genomic occupancy, transcription-factor interactions, and effects on local chromatin and promoter activity (Barski et al., 2010; Ferrigno et al., 2001; Gerber et al., 2020; Yague-Sanz et al., 2023). Pol II binds to or near the promoters of many Pol III-transcribed housekeeping genes to support their expression (Listerman et al., 2007; Raha et al., 2010). Several Pol II-associated transcription factors and downstream RNA cleavage and polyadenylation components can also be recruited to these Pol III genes (Lari et al., 2025; Raha et al., 2010). More recently, the reciprocal relationship has been demonstrated, revealing that Pol III broadly occupies the promoters of Pol II-transcribed genes and can impact their expression (Jiang et al., 2022; K C et al., 2024). This suggests that Pol III occupancy at protein-coding gene promoters has regulatory potential, which we hypothesized could be relevant under conditions of pathogenic stress.

Here, we investigated the role of Pol III in shaping the cellular transcriptome in both resting and virally infected cells. We used the murine model gammaherpesvirus MHV68, which is related to the human oncogenic herpesviruses Kaposi’s sarcoma-associated herpesvirus (KSHV) and Epstein-Barr virus (EBV), and robustly activates Pol III transcription of the murine B1 and B2 SINE families (Karijolich et al., 2017; Lari et al., 2025). We found that Pol III performs two distinct functions at Pol II promoters. In resting cells, it supports chromatin accessibility and basal priming of innate immune genes. During infection with multiple viruses, including MHV68, herpes simplex virus-1 (HSV-1), and influenza A virus (IAV), it restrains the magnitude of inflammatory gene induction. Pol III acts at Pol II promoters with specific architecture and may thus compete with Pol II for shared transcriptional machinery. Together, these findings establish Pol III as a direct-acting regulator of antiviral gene expression.

## RESULTS

### RNA polymerase III is redistributed from protein-coding gene promoters to SINE retro-transposons during infection

To first assess whether Pol III occupancy at protein-coding genes is altered during viral infection, we examined our published Polr3A chromatin immunoprecipitation sequencing (ChIP-seq) dataset from mock-treated and MHV68-infected NIH3T3 fibroblasts (Lari et al., 2025). Consistent with reports in human cancer cells and mouse embryonic stem cells (Jiang et al., 2022; K C et al., 2024), Polr3A, the major catalytic subunit of the Pol III transcription complex, was broadly enriched at the promoters of protein-coding genes in uninfected cells (**Fig. 1A**). Polr3A consensus peaks were tightly focused at annotated transcription start sites (TSSs), with 50.6% falling within ± 100 bp of a protein-coding gene transcription start site, of which only 0.6% overlapped tRNA genes and 3.8% overlapped infection inducible B2 SINE elements (**Supplemental Fig. 1A**). Thus, Pol III occupies core Pol II promoter regions that are distinct from its binding at canonical Pol III-transcribed elements.

**Figure 1.**
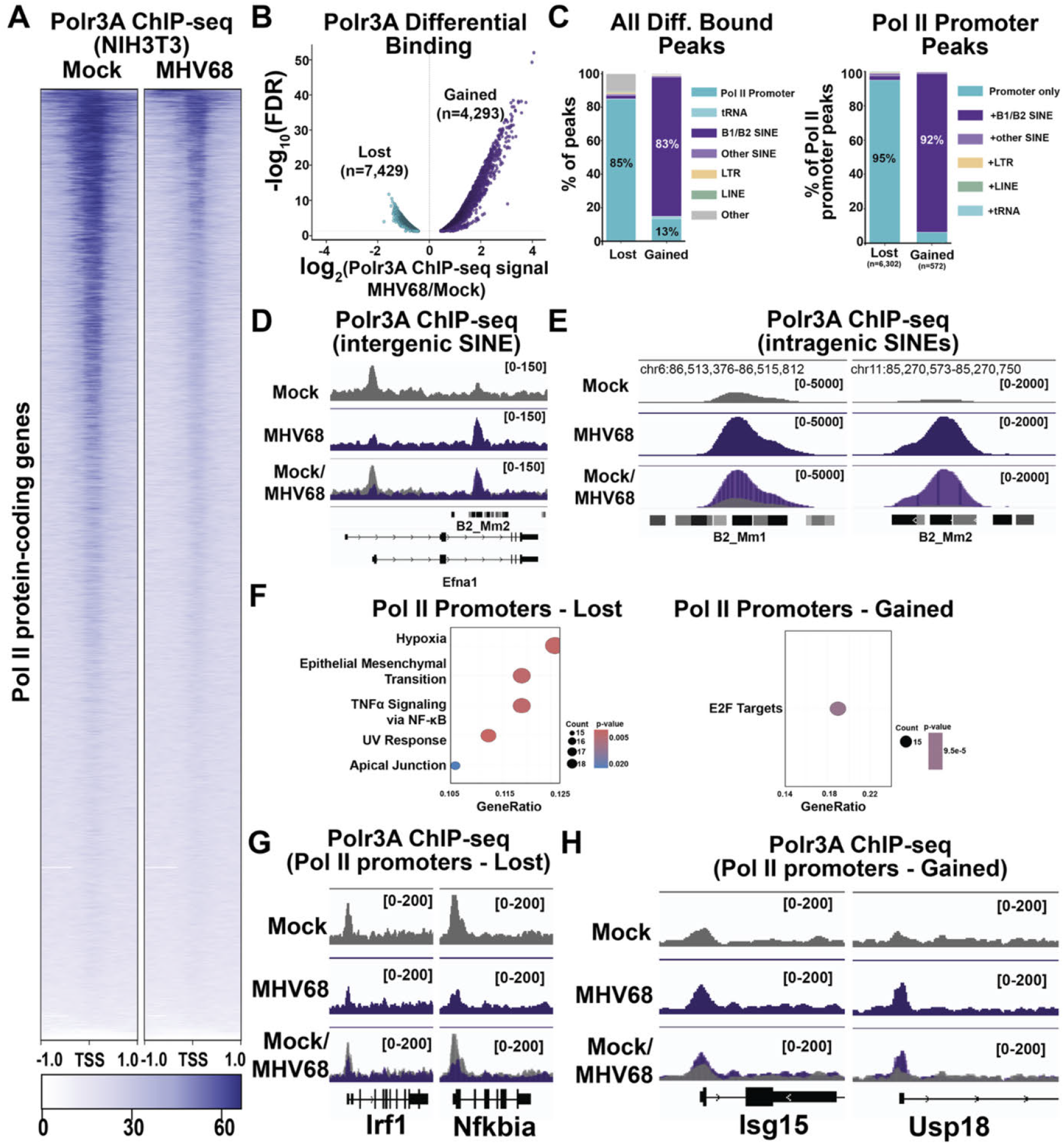
RNA Polymerase III occupancy is redistributed from protein-coding gene promoters to infection-inducible SINE loci during MHV68 infection. **A**. Heatmap displaying Polr3A ChlP-seq signal across protein-coding genes in mock-treated or MHV68-infected NIH3T3 cells (n=21,962).. ChlP-seq signal was plotted as a histogram with 10-bp bins from −1 to +1 kbp around the transcription start site (TSS), with 20 bins/gene. **B**. Volcano plot of Polr3A differential binding analysis comparing MHV68-infected to mock cells. Significant peaks (FDR < 0.05) are colored by direction: lost (n = 7,429; teal) and gained (n = 4,293; purple). Dashed horizontal line indicates FDR = 0.05. **C. left**: Genomic-context classification of significantly differentially bound Polr3A peaks (FDR < 0.05). **Right**: Sub-classification of Polr3A peaks within the Pol II promoter category by repeat-element overlap status within the promoter region. **D**. Polr3A ChlP-seq coverage across a representative B2 SINE gene that is embedded within the intron of a proteincoding gene. **E**. PoIr3A ChlP-seq coverage across a representative intergenic B2 SINE gene. F. MSigDB Hallmark enrichment of high-confidence differentially bound Pol II promoter peak sets (FDR < 0.05, logaFC > 1). **Left:** SINE-free promoters that lose Polr3A occupancy (n = 410 genes). **Right:** SINE-overlapping promoters that gain Polr3A occupancy (n - 309 genes). Dot size indicates gene count, and dot color indicates the Benjamini-Hochberg (BH)-adjusted p-value. **G**. Representative Polr3A ChlP-seq tracks at two inflammatory transcription factor genes (*Irf1, Nfkbia*) that lose TSS-localizcd Polr3A signal during MH V68 infection. **H**. Representative Polr3A ChlP-seq tracks at two interferon-stimulated genes (*Isg15, Uspl8*) that gain Polr3A occupancy at their promoter regions during MHV68 infection.

MHV68 infection led to a marked, genome-wide redistribution of Pol III (**Fig. 1A**). Differential binding analysis identified 7,429 Polr3A peaks that lost signal and 4,293 peaks that gained signal during infection (**Fig. 1B**). Classification of these differentially bound peaks by genomic context revealed two opposite patterns: lost peaks were predominantly located at Pol II protein-coding gene promoters (85%), while gained peaks were predominantly at B1/B2 SINE retrotransposons (83%) (**Fig. 1C, left**). Representative tracks at intergenic (**Fig. 1D**) and intragenic (**Fig. 1E**) B2 SINE loci illustrate this gain of Pol III at retrotransposons during infection. While the total integrated Polr3A ChIP-seq signal across all peaks was largely unchanged between mock and MHV68 conditions (**Supplemental Fig. 1B, left**), the per-peak signal showed substantial reorganization consistent with Pol III being redistributed across chromatin rather than evicted or degraded during infection (**Supplemental Fig. 1B, right**). Consistent with this redistribution, protein-coding gene promoters showed significantly greater loss of Polr3A signal during infection than tRNA/SINE loci, indicating that Pol III is preferentially retained at canonical targets while vacating protein-coding promoters (**Supplemental Fig. 1C**). Protein levels of four core Pol III subunits also remained stable across a time course of MHV68 lytic replication (**Supplemental Fig. 1D**). A finite pool of Pol III is therefore dynamically partitioned between Pol II promoters and SINE retrotransposons during infection.

To further resolve the lost and gained populations of Polr3A peaks at protein-coding gene promoters, we examined whether individual peaks localized directly to the annotated TSSs or rather with a promoter-proximal SINE element. Strikingly, 95% of lost-Polr3A promoter peaks occupied SINE-free TSS regions, while 92.3% of gained-Polr3A promoter peaks overlapped a B1/B2 SINE within the promoter region (**Fig. 1C, right**).

Functional enrichment analysis of the two populations revealed that protein-coding gene promoters that lost Polr3A binding during MHV68 infection were enriched for TNF*α* signaling via NF-*κ*B, hypoxia response, and epithelial-mesenchymal transition (**Fig. 1F, left**). This population also included inflammatory transcription factors such as Irf1, Irf3, Nfkbia, and Fosl2. Representative tracks at two promoters illustrate the direct TSS-localized loss of Polr3A during infection (**Fig. 1G**). Conversely, promoter regions that gained Polr3A at adjacent B1/B2 SINE elements were enriched for E2F target genes (**Fig. 1F, right**), a class of cell cycle and proliferation regulators that gammaherpesviruses exploit to establish replication-permissive cellular states (Hume and Kalejta, 2009). Promoter regions that gained Polr3A also included canonical interferon-stimulated genes (ISGs), including Isg15, Usp18, Tyk2, Ifnar2, and Oasl2 (**Fig. 1H**).

Together, these findings indicate that Pol III engages two distinct populations of host gene promoters that are reciprocally regulated during MHV68 infection: a dominant population of Polr3A binding at the TSSs of inflammatory regulators, which loses signal during infection, and a smaller population of SINE-mediated Polr3A recruitment to E2F target genes and a subset of ISGs.

### RNA polymerase III supports basal innate immune and inflammatory gene expression

We next defined the functional consequences of Pol III occupancy at Pol II promoters using primary murine bone marrow-derived macrophages (BMDMs), an innate immune cell model relevant to MHV68 infection in vivo (Herskowitz Jeremy et al., 2008; Weck et al., 1999). To prevent Pol III recruitment to these promoters, we depleted Brf1 (**Fig. 2A** and **Supplemental Fig. 2A**), a component of the Pol III transcription factor TFI-IIB complex enriched at Pol III-occupied promoters of protein-coding genes in both human and murine cells (Carrière et al., 2011; K C et al., 2024). Consistent with preferential Brf1 recruitment at protein-coding loci, our analyses of published ChIP-seq data (Carrière et al., 2011) showed little to no occupancy of the Brf1 paralog, Brf2 (**Supplemental Fig. 2B-C**), which instead directs Pol III recruitment to type III (e.g., U6, 7SK, Y RNAs) promoters (Schramm and Hernandez, 2002).

**Figure 2.**
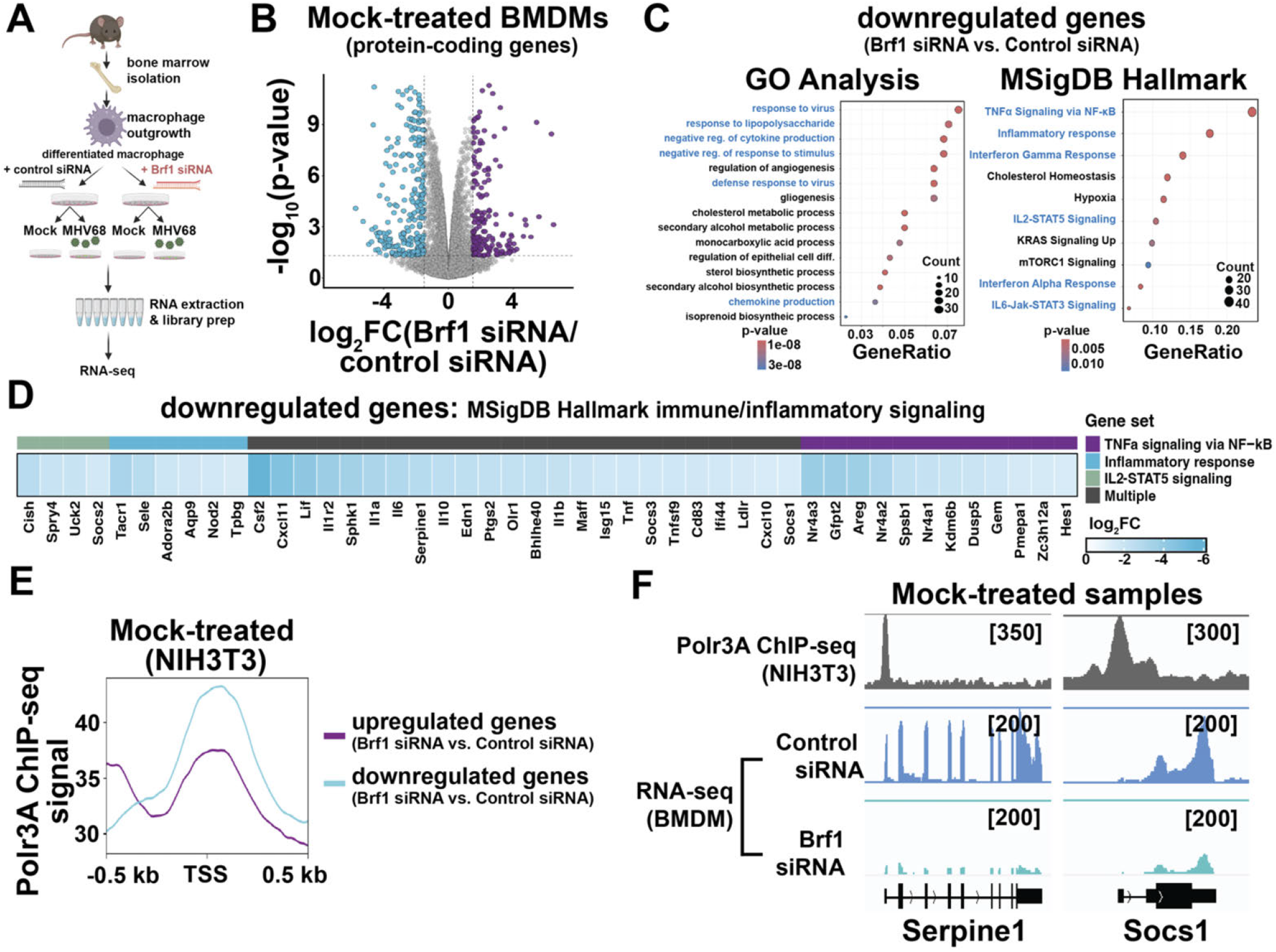
RNA Polymerase III activity impacts innate immune and inflammatory gene expression in resting primary BMDMs. **A**. Schematic showing the workflow of RNA-seq experiments in which primary BMDMs transfected with control non-targeting or Brf 1-targeting siRNAs were mock-treated or infected with MHV68 at an MOI of 5 in triplicate. **B**. Differential expression analysis at proteincoding genes (n= 14,416) from RNA-seq was plotted as fold change (FC) versus adjusted p-value. A p-value threshold of <0.05 was used to identify genes that were significantly downregulated (log_2_FC ≤ −1.5, cyan, n=255) or upregulated (log_2_FC ≥ 1.5, purple, n=175) when comparing primary BMDMs transfected with Brfl targeting siRNAs to those transfected with control non-targeting siRNA under mock conditions. **C**. Gene Ontology Biological Process enrichment (left) and MSigDB Hallmark gene-set enrichment (right) of Brfl-dependent downregulated protein-coding genes in BMDMs. Dot size indicates gene count, and dot color indicates the Benjamini-Hochberg (BH)-adjusted p-value. **D**. Heatmap of logz fold change (Brfl _si_RNA vs. control siRNA, mock-treated) for significantly downregulated genes (adjusted P < 0.05, log2FC ≤ −1.5) that overlap one or more MSigDB Hallmark gene sets associated with inflammatory and cytokine signaling: TNF α signaling via NF-_k_B, inflammatory response, interferon a response, interferon y response, IL2-STAT5 signaling, and IL6-JAK-STAT3 signaling. Genes are annotated by gene-set membership (color bar); genes belonging to multiple sets are labeled “Multiple.” **E**. Metagene plot displaying Polr3A ChlP-seq signal across differentially expressed protein-coding gene sets from (B). The ChlP-seq signal was plotted as a histogram with 10-bp bins from −0.5 to +0.5 kb around the transcription start site (TSS), with 20 bins/genc. **F**. Polr3A ChlP-seq and RNA-seq coverage across representative protein-coding genes from the downregulated gene set from (B).

We measured how Pol III occupancy influences gene expression in BMDMs using RNA-seq with spike-in controls (**Supplemental Fig. 2D**). To avoid confounding signal from intragenic Pol III-transcribed SINE elements, RNA-seq analyses were restricted to annotated exonic regions, which generally exclude embedded SINE sequences (Zhang et al., 2011). In uninfected BMDMs, Brf1 depletion caused differential expression of approximately 400 protein-coding genes (**Fig. 2B** and **Supplemental Table 1**). Strikingly, the downregulated gene set was strongly enriched for innate immune and inflammatory pathways by both MSigDB Hallmark and Gene (GO) analyses (**Fig. 2C**). The top Hallmark categories included TNF*α* signaling via NF-*κ*B, inflammatory response, interferon gamma response, interferon alpha response, and IL2-STAT5 and IL6-JAK-STAT3 signaling. GO analysis confirmed this signature. Individual Brf1-dependent inflammatory genes included canonical innate immune effectors such as Tnf, Il1a, Il1b, Il6, Cd83, Serpine1, and Socs1 (**Supplemental Table 1** and **Fig. 2D**). Many of the differentially expressed transcripts exhibited relatively low expression levels (**Supplemental Fig. 2E**), consistent with a role for Pol III in maintaining a basal state of low-level innate immune gene expression in uninfected BMDMs. In contrast, the upregulated set was much smaller and was enriched for the complement pathway and for cellular migration and chemotaxis programs by GO analysis (**Supplemental Fig. 2F**), indicating that Brf1 loss does not broadly de-repress host genes in resting BMDMs but largeley dampens the innate immune effector program.

The Brf1-dependent downregulated gene set showed substantially higher Polr3A occupancy at their promoters in uninfected cells than the upregulated gene set (**Fig. 2E-F**), establishing a connection between Pol III binding and basal inflammatory gene expression. Notably, TNF*α*/NF-*κ*B pathway enrichment within the Brf1-dependent gene set in BMDMs mirrored the enrichment observed at Polr3A-occupied promoters in fibroblasts (**Fig. 1F**), supporting a model in which Pol III occupancy at innate immune promoters is a conserved, relevant feature across cell types.

### RNA polymerase III activity restrains inflammatory gene expression during infection

We next asked how Pol III shapes gene expression during infection, when it moves from Pol II promoters to SINEs. RNA-seq of MHV68-infected primary BMDMs (48 hpi) following Brf1 depletion identified 373 upregulated and 381 downregulated protein-coding genes (**Fig. 3A, Supplementary Table 2**). While the downregulated set was very lowly expressed and showed no coherent functional enrichment (**Supplemental Fig. 3A-B**), the upregulated set was enriched for inflammatory genes. These included TNF*α* signaling via NF-*κ*B, inflammatory response, complement, IL2-STAT5 signaling, IL6-JAK-STAT3 signaling, and p53 pathway by MSigDB Hallmark analysis, and regulation of inflammatory response by GO Biological Process analysis (**Fig. 3B-C**). Notably, Polr3A was differentially lost from the promoters of this upregulated gene set during infection (**Fig. 3D-E**), further suggesting that Pol III occupancy restrains Pol II activity at induced promoters.

**Figure 3.**
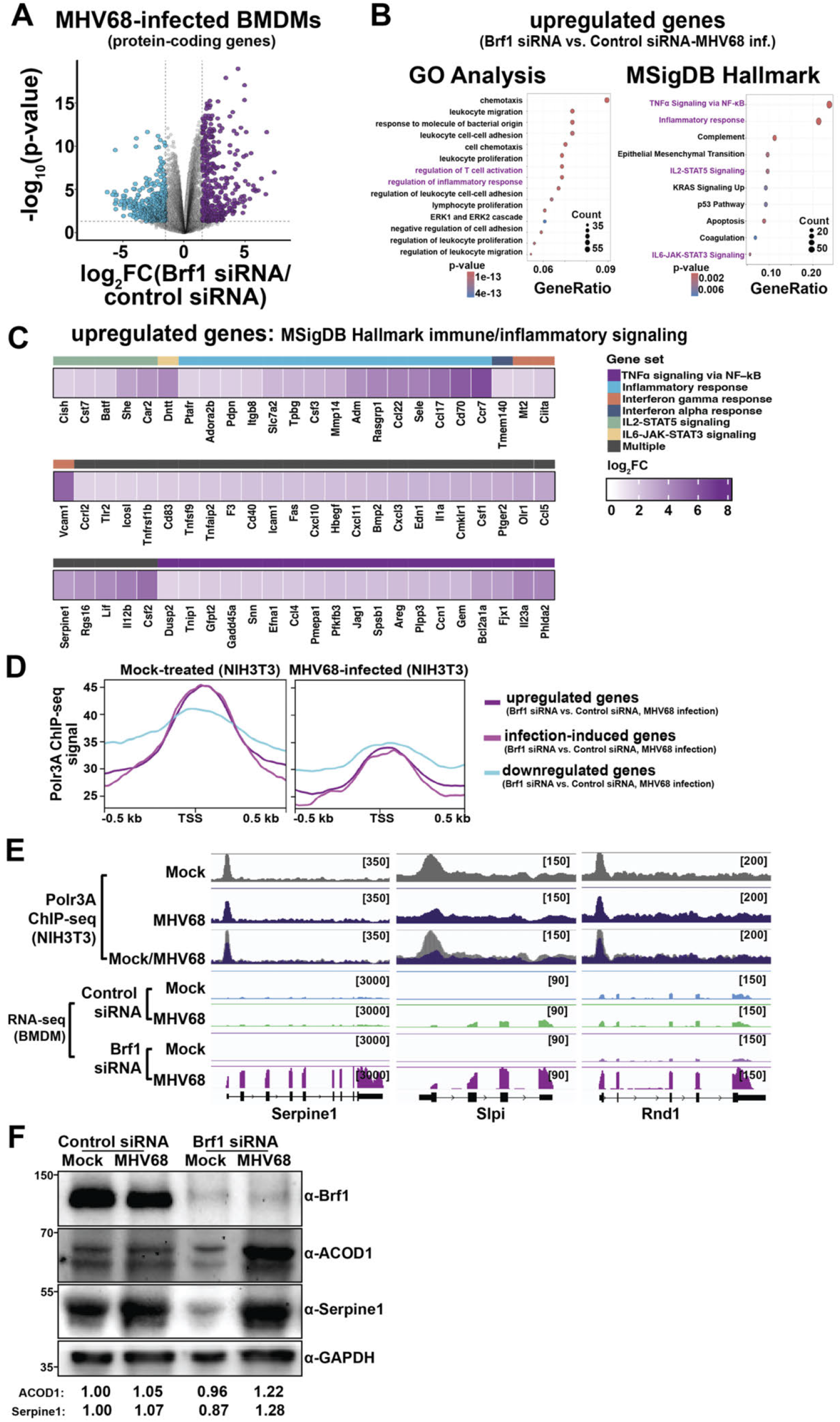
RNA Polymerase Ill activity restricts innate immune and inflammatory gene expression in MHV68-infectcd primary BMDMs. **A**. Differential expression analysis at protein-coding genes (n=13,367) from RNA-seq was plotted as fold change (FC) versus adjusted p-value. A p-value threshold of <0.05 was used to identify genes that were significantly downregulated (log_2_FC ≤ −1.5, cyan, n=381) or upregulated (log_2_FC ≥ 1.5, purple, n=373) when comparing MHV68-infected primary BMDMs transfected with Brfl targeting siRNAs to those transfected with control non-targeting siRNA. **B**. Gene Ontology Biological Process enrichment and MSigDB Hallmark gene-set enrichment of Brfl-dependent upregulated protein-coding genes in MHV68-infected BMDMs. Dot size indicates gene count, and dot color indicates the Benjamini-Hochberg (BH)-adjusted p-value. **C**. Heatmap of logz fold change (Brfl siRNA vs. control siRNA, 48 hpi) for significantly upregulated genes (adjusted P < 0.05, log_2_FC ≥ 1.5) that overlap one or more MSigDB Hallmark gene sets associated with inflammatory and cytokine signaling: TNFa signaling via NF-kB, inflammatory response, interferon a response, interferon γ response, IL2-STAT5 signaling, and IL6-JAK-STAT3 signaling. Genes are annotated by gene-set membership (color bar); genes belonging to multiple sets are labeled “Multiple”. **D**. Metagene plot displaying Polr3A ChlP-seq signal across differentially expressed protein-coding gene sets from (A). This includes upregulated genes (purple), downregulated genes (cyan), and upregulated, infection-induced genes (magenta). ChlP-seq signal was plotted as a histogram with 10 bp bins from −0.5 to +0.5 kbp around the transcription start site (TSS), with 20 bins/gene. **E**. Polr3A ChlP-seq and RNA-seq coverage across a select protein-coding genes from the upregulated and infection-induced gene sets from (A). **F**. Primary BMDMs transfected with control non-targeting, or (Brf1-targeting siRNAs were mock-treated or infected with MHV68 at an MOI of 5. At 48 hpi), cells were harvested and lysed to extract total protein, which was then analyzed by Western blotting using antibodies against Brf1, Acodl, Serpinel, and GAPDH (loading control). The relative protein levels of Acodl and Serpinel were measured as the ratio of their mean integrated intensities to GAPDH levels, normalized to mock-treated controls treated with control siRNA.

Although the genes upregulated upon Brf1 depletion generally exhibited lower-than-average expression levels, some were highly expressed, with levels consistent with immune activation during infection (**Supplemental Fig. 3B**). We confirmed for two of these upregulated, highly expressed immune-responsive factors (Acod1 and Serpine1) that the Pol III-dependent transcript-level changes observed in uninfected and infected cells reflected protein-level changes (**Fig. 3F**). Brf1 depletion did not alter MHV68 gene expression or viral DNA replication (**Supplemental Fig. 4A-B**), establishing that the inflammatory phenotype is not driven by increased viral levels. Thus, Pol III activity has opposing effects on inflammatory pathway genes in resting versus infected cells: it promotes their basal expression in uninfected BMDMs but restrains their expression during MHV68 infection, perhaps helping prevent excessive inflammation.

### RNA polymerase III activity shapes mRNA isoform diversity at innate immune, inflammatory, and stress-response genes

Our observations suggested that Pol III shapes the magnitude of transcriptional output at infection-responsive promoters. We next asked whether Pol III also impacts the specific mRNA isoforms produced, since alternative splicing, transcription start/end site selection, and intron retention are critical regulatory layers of inflammatory gene expression (Boudreault et al., 2016; Ivanov and Anderson, 2013; Jia et al., 2017; Pai et al., 2016). To define full-length transcript structures, we performed PacBio Iso-Seq and isoform characterization analysis (SQANTI - (Tardaguila et al., 2018)) on biological duplicates of BMDMs treated with control or Brf1-targeting siRNAs in both mock and MHV68-infected conditions, which were matched the results to our short-read RNA-seq datasets.

Brf1 depletion led to widespread isoform remodeling in both uninfected and MHV68-infected BMDMs (**Fig. 4A-B** and **Supplemental Tables 3-4**). In uninfected cells, 550 isoforms corresponding to 439 protein-coding genes were differentially expressed upon Brf1 depletion. In MHV68-infected cells, 224 isoforms were upregulated and 129 downregulated. In both contexts, approximately half of the isoform-level changes occurred independently of gene-level expression changes, indicating that Pol III activity regulates transcript structure at a substantial set of genes that do not show overt expression differences (**Fig. 4C-D**). The isoform-level changes encompassed intron retention, alternative splicing, and shifts in both transcription start and end sites (**Fig. 4E**). Notably, in MHV68-infected cells, the downregulated isoforms were highly enriched for intron retention events (**Fig. 4F**). The Rbm39 locus provides a representative example of an isoform-level change that was not detected by gene-level differential expression analysis. Although total Rbm39 expression was unchanged following Brf1 depletion, RNA-seq coverage was altered across a retained intronic region (**Fig. 4G**). These findings suggest that Pol III activity can influence the relative abundance of intron-retaining transcripts during infection.

**Figure 4.**
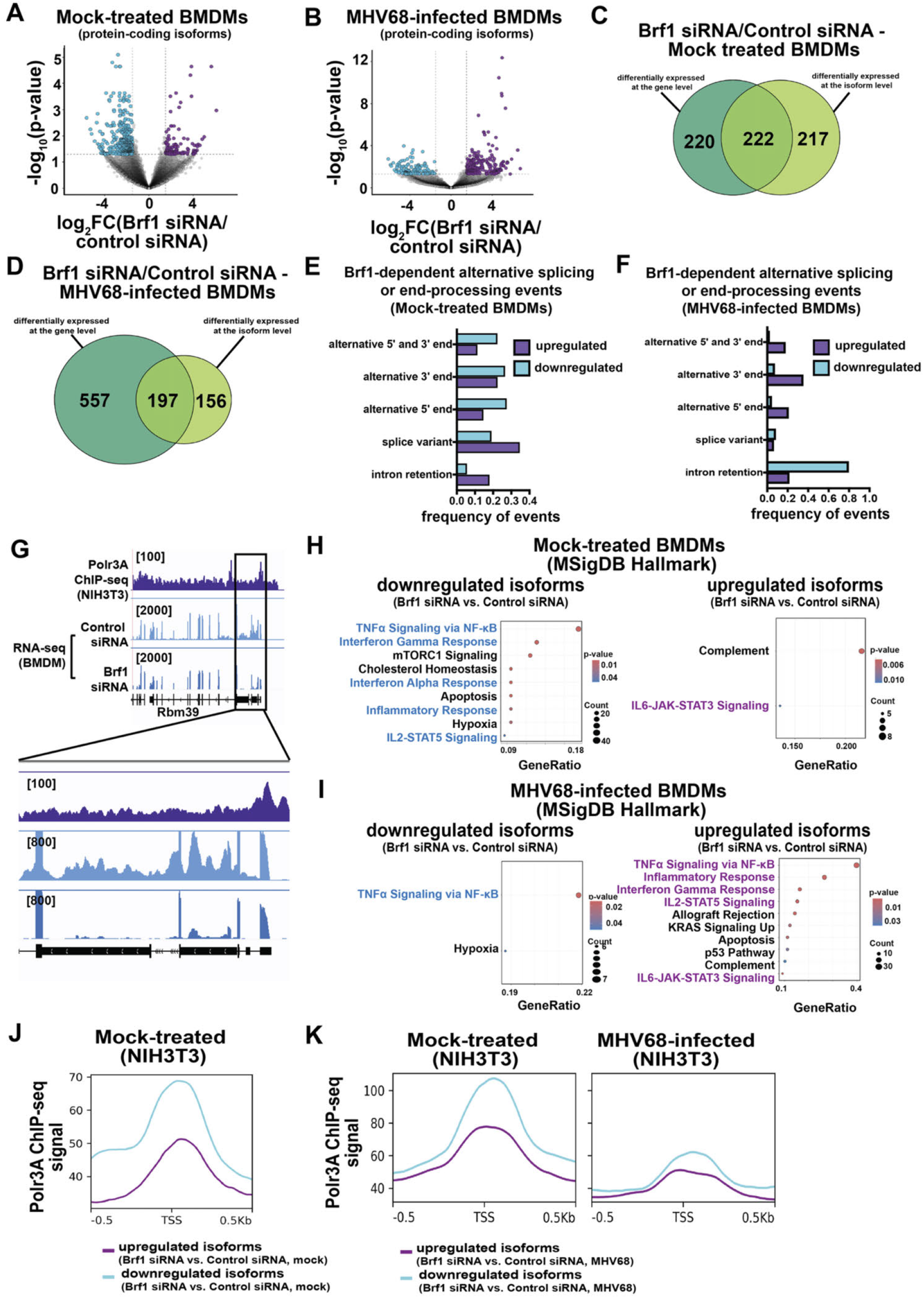
RNA Polymerase III activity influences mRNA isoform diversity in primary BMDMs. **A**. Differential isoform expression analysis at protein-coding genes (n= 14416) from long-read RNA-seq was plotted as fold change (FC) versus adjusted p-value. A p-value threshold of <0.05 was used to identify isoforms that were significantly downregulated (log_2_FC ≤ −1.5, cyan) or upregulated (log_2_FC ≥1.5, purple) when comparing primary BMDMs transfected with Brfl targeting siRNAs to those transfected with control non-targeting siRNA under mock conditions. **B**. Differential isoform expression analysis at protein-coding isoforms (n= 13367) was plotted as fold change (FC) versus adjusted p-value. A p-value threshold of <0.05 was used to identify isoforms that were significantly downregulated (log_2_FC ≤ −1.5, cyan) or upregulated (log_2_FC ≥ 1.5, purple) when comparing MHV68-infected primary BMDMs transfected with Brfl targeting siRNAs to those transfected with control non-targeting siRNA. **C**. Venn diagram showing the number of genes that overlap between differential expression at the gene level (Fig. 2B) or isoform level from (A) when comparing primary BMDMs transfected with Brfl targeting siRNAs to those transfected with control non-targeting siRNA under mock conditions. **D**. Venn diagram showing the number of genes that overlap between differential expression at the gene level (Fig. 2B) or isoform level from (A) when comparing MHV68-infected primary BMDMs transfected with Brfl targeting siRNAs to those transfected with control non-targeting siRNA. **E**. Graph shows the frequency of various splicing and transcription start and end site usage events associated with differentially expressed isoforms from (A) as analyzed by SQANTI (Tardaguila et al., 2018). **F**. Graph shows the frequency of various splicing and transcription start and end site usage events associated with differentially expressed isoforms from (B) as analyzed by SQANTI (Tardaguila et al., 2018). **G**. Polr3A ChlP-seq and RNA-seq coverage across a representative protein-coding gene showing a retained intron from the downregulated isoform set from (A). **H**. MSigDB Hallmark gene-set enrichment of differentially expressed isoforms from PacBio Iso-Seq in mock-treated BMDMs (Brfl siRNA/control siRNA). Left: isoforms downregulated upon Brfl depletion. Right: isoforms upregulated upon Brfl depletion. Dot size indicates gene count, and dot color indicates the Benjamini-Hochberg (BH)-adjustcd p-value. **I**. MSigDB Hallmark gene-set enrichment of differentially expressed isoforms in MHV68-infected BMDMs at 48 hpi (Brfl siRNA/control siRNA); isoform analysis performed as in (B). Left: downregulated isoforms. Right: upregulated isoforms. Dot size indicates gene count, and dot color indicates the Benjamini-Hochberg (BH)-adjusted p-value. **J**. Metagene plots displaying Polr3A ChlP-seq signal across differentially expressed protein-coding isoform sets from (A). This includes upregulated (purple) and downregulated (cyan) isoform sets, comparing mock-treated primary BMDMs transfected with Brfl-targeting siRNAs to those transfected with control non-targeting siRNAs. The ChlP-seq signal was plotted as a histogram with 10-bp bins from −0.5 to +0.5 kb around the transcription start site (TSS), with 20 bins/gene. **K**. Mctagcnc plots displaying Polr3A ChlP-seq signal across differentially expressed protein-coding isoform sets from (B). This includes upregulated (purple) and downregulated (cyan) isoform sets, comparing MHV68-infected primary BMDMs transfected with Brfl-targeting siRNAs to those transfected with control non-targeting siRNAs. The ChlP-seq signal was plotted as a histogram with 10-bp bins from −0.5 to +0.5 kb around the transcription start site (TSS), with 20 bins/gene.

MSigDB Hallmark enrichment of the differentially regulated isoforms revealed the same context-dependent inflammatory signature observed at the gene level. In uninfected cells, downregulated isoforms were enriched for, e.g., TNF*α* signaling via NF-*κ*B, interferon alpha and gamma, and inflammatory responses, as well as IL2-STAT5 signaling (**Fig. 4H**). In MHV68-infected cells, the upregulated isoforms were strongly enriched for the same canonical inflammatory programs, including TNF*α*/NF-*κ*B, inflammatory response, interferon gamma response, and IL2-STAT5 signaling (**Fig. 4I**). Thus, Pol III influences both expression levels and isoform production for inflammatory pathway genes, with opposing outcomes in uninfected versus infected cells.

Consistent with a direct role for Pol III occupancy in shaping isoform-level expression, isoforms whose expression decreased upon Brf1 depletion in both uninfected and infected conditions arose from promoters with substantially higher Polr3A occupancy than those whose expression increased (**Fig. 4J-K**). By contrast, the upregulated isoforms during MHV68 infection that were further amplified by Brf1 depletion exhibited only modest Polr3A occupancy at their promoters in both mock and MHV68 conditions (**Fig. 4K**). This pattern indicated that the large infection-induced redistribution of Pol III away from Pol II promoters primarily affected gene-level expression, while isoform-level changes may additionally have arisen from local Pol III-dependent influences on Pol II elongation, splicing, and transcription termination. Thus, Pol III occupancy also shapes the structure of the resulting mRNAs, with the same Pol III-dependent inflammatory genes producing distinct isoforms in different cellular states.

### RNA polymerase III maintains chromatin accessibility at RNA polymerase II promoters

To address how Pol III regulates expression of innate immune genes, we examined whether it influences the chromatin environment at these promoters. We performed ATAC-seq with spike-in normalization in NIH3T3 fibroblasts depleted of Brf1, with and without MHV68 infection, to match the cell type of our ChIP-seq dataset. In uninfected cells, Brf1 depletion reduced chromatin accessibility at Pol II protein-coding gene promoters in a progressive, Polr3A dose-dependent manner (**Fig. 5A-B**). Stratifying 20,000 protein-coding gene promoters by their Polr3A occupancy, the top quintile of Polr3A-occupied promoters (Q5; mean spike-in-normalized signal = 95) showed the largest accessibility decrease (6% absolute reduction vs lowest quintile; **Fig. 5B**). Absolute mean ATAC signal per quintile confirmed that the high-Pol-III accessibility loss is genuine and not driven by ratio-based normalization (**Supplemental Fig. 5A**). Furthermore, the Brf1-dependent inflammatory gene promoters (from **Fig. 3A**) showed a substantially larger ATAC-seq signal decrease in Brf1-depleted fibroblasts than promoters of unchanged genes (**Fig. 5C**). These data indicate that Pol III maintains a permissive chromatin state at the protein-coding gene promoters it occupies and supports their basal expression. The Pol III-dependent chromatin licensing function was also observed at canonical Brf1-dependent genes (tRNAs, SINEs) in both uninfected and infected cells, with MHV68 infection itself dramatically remodeling chromatin accessibility at these targets (**Supplemental Fig. 5B-C**).

**Figure 5.**
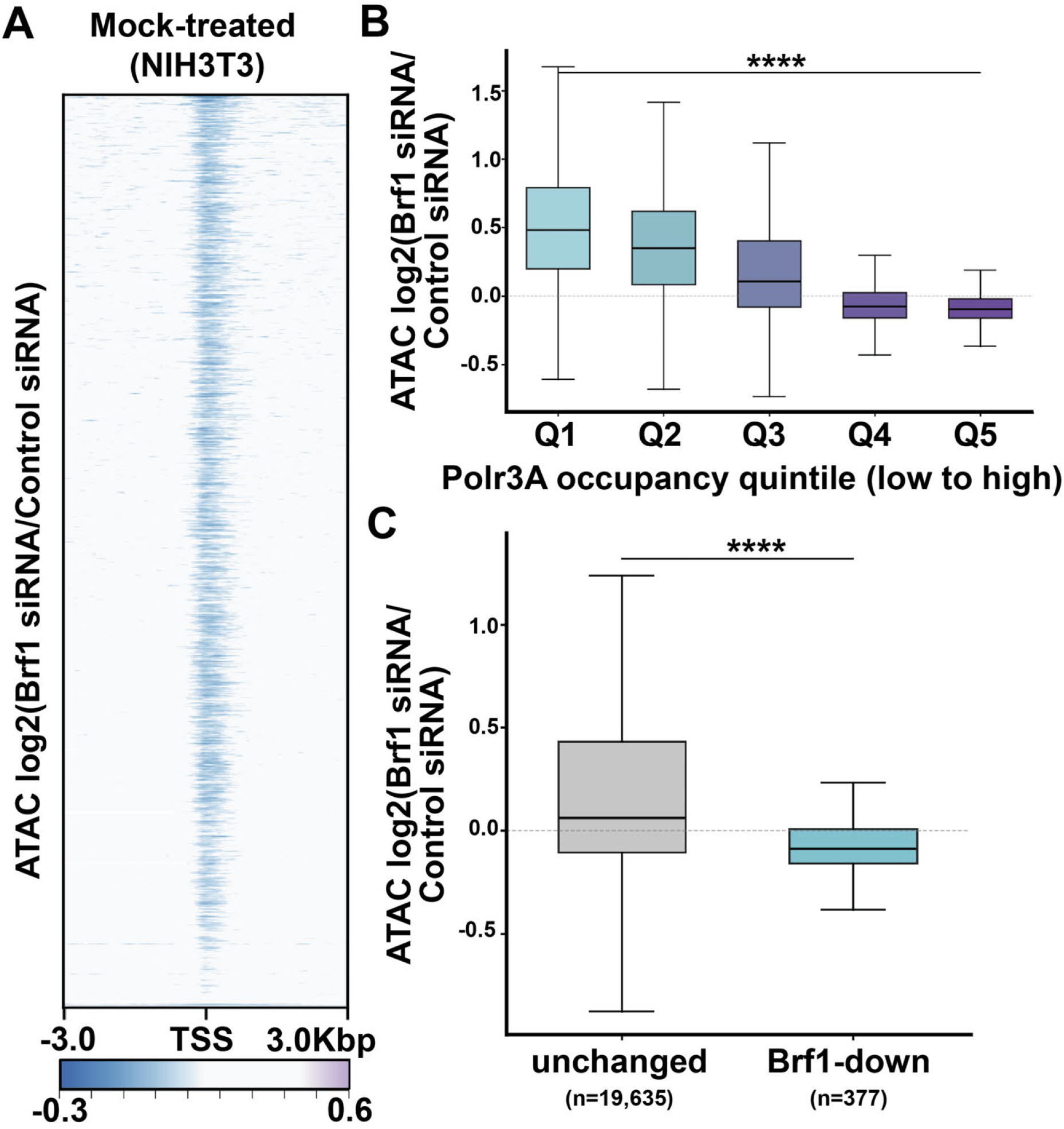
RNA Polymerase III maintains chromatin accessibility at the Pol II protein-coding gene promoters. **A**. Heatmap of ATAC-seq log_2_(Brfl siRNA/control siRNA) at protein-coding gene transcription start sites (TSSs; ± 3 kb) in mock-treated NIH3T3 fibroblasts (n = 20,012 protein-coding genes). Color scale: log2(Brfl siRNA/control siRNA); purple, accessibility gain; teal, accessibility loss; white, no change (clipped at ± 0.6). **B.** ATAC-seq log_2_(Brfl siRNA/control siRNA) at protein-coding gene promoters (TSS ±500 bp), stratified by Polr3A occupancy quintile in uninfected NIH3T3 fibroblasts. Each box represents ~ 4,000 promoters. Boxes show median and interquartile range; whiskers show 1.5 x IQR; outliers omitted for clarity. Statistical comparison of Q5 vs QI distributions: two-sided Mann-Whitney U test, ^* * * *^ P < 10^−4^. **C.** ATAC-seq log_2_(Brfl siRNA/control siRNA) at promoters of Brfl-dependent downregulated inflammatory genes (n = 377 mapped of 410 BMDM Brfl-KD downregulated genes from Fig. 2) compared to all unchanged genes (n = 19,635). Boxes show median and interquartile range; whiskers show 1.5 x IQR; outliers omitted for clarity. Statistical comparison: two-sided Mann-Whitney U test, ^* * * *^ P < 10^−4^.

In contrast to uninfected cells, Brf1 depletion during MHV68 infection led to little change in chromatin accessibility at Polr3A-occupied Pol II protein-coding gene promoters (**Supplemental Fig. 6A**). The dose-response relationship across Polr3A quintiles was also substantially attenuated (**Supplemental Fig. 6B**), and the Pol III–restrained inflammatory genes identified by RNA-seq in infection conditions (**Fig. 3A**) showed no measurable accessibility difference from unchanged genes (**Supplemental Fig. 6C**). The amplification of inflammatory transcription upon Brf1 depletion during infection (**Fig. 3**), therefore, likely reflects a chromatin-independent mechanism.

### RNA polymerase III also regulates inflammatory gene expression during human DNA and RNA virus infections

To determine whether the regulation of infection-responsive genes by Pol III extended to human virus infections, we performed RNA-seq in A549 lung epithelial cells infected with herpes simplex virus-1 (HSV-1) and influenza A virus (IAV), with and without Brf1 depletion. Like MHV68, these viruses induce Pol III-driven SINE gene expression (Dremel et al., 2022; Shen et al., 2022). In both cases, Brf1 depletion led to widespread differential protein-coding gene expression (HSV-1: 532 genes upregulated; IAV: 708 genes upregulated; **Fig. 6A-B** and **Supplemental Tables 5-6**).

**Figure 6.**
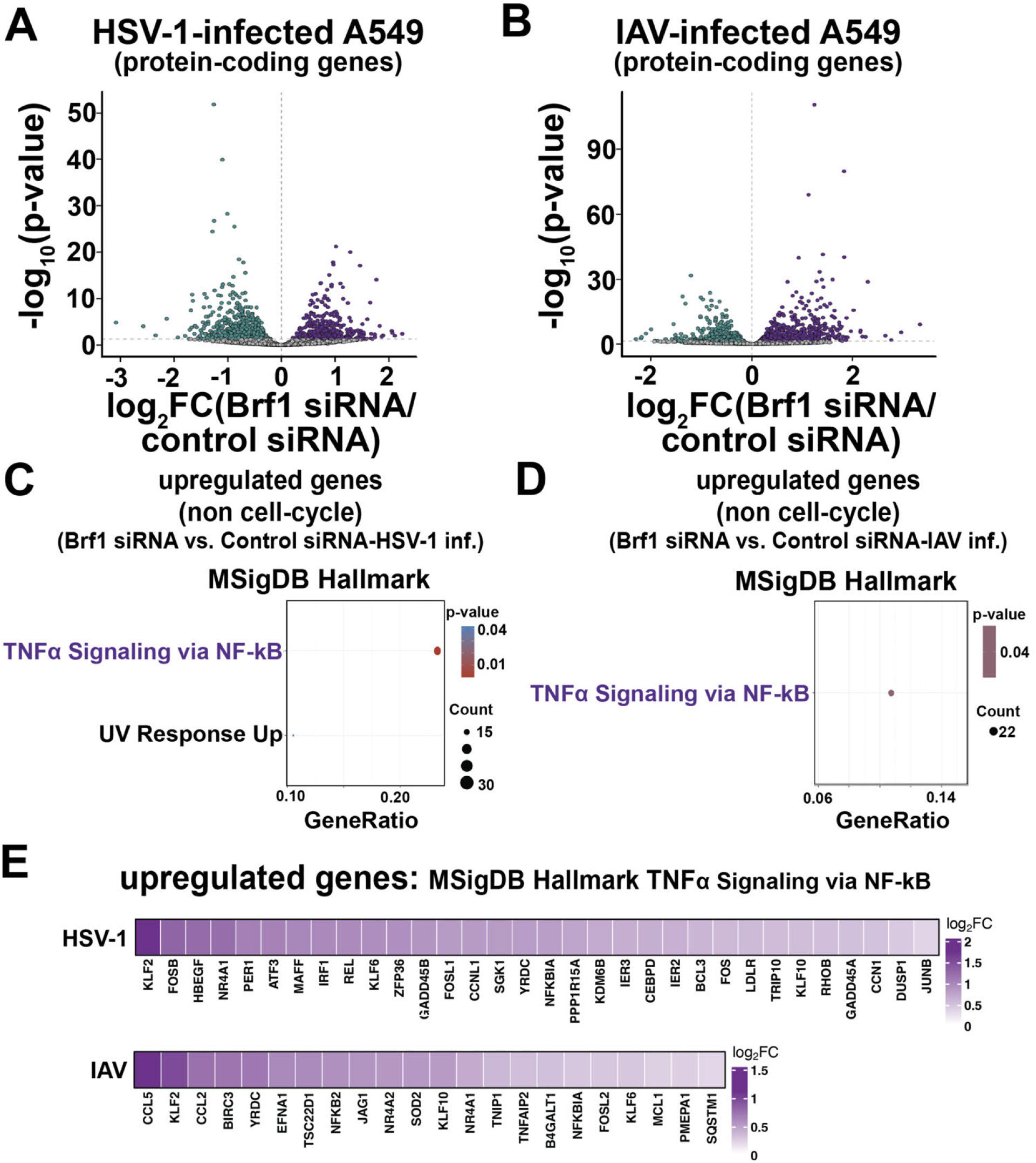
Brfl depletion derepresses host genes during HSV-1 and 1AV infection of human A549 cells. **A**. Volcano plot of differential gene expression (protein-coding genes) in HSV-1-infected A54 9 cells transfected with Brfl-targeting siRNA versus control siRNA. The x-axis shows log_2_ fold change (Brfl siRNA / control siRNA) and the x-axis shows-logic _10_(p-value). Magenta points indicate genes significantly upregulated and cyan points indicate genes significantly downregulated upon Brfl depletion (p-value < 0.05; grey points are not significantly changed. **B**. MSigDB Hallmark gene-set enrichment for the genes upregulated upon Brfl depletion in HSV-1-infected cells (Brfl siRNA vs. control siRNA). The dot position on the x-axis indicates gene ratio, the dot size indicates the number of genes in each set (Count), and the dot color indicates enrichment p-value. **C**. Volcano plot of differential gene expression (proteincoding genes) in lAV-infcctcd A549 cells transfected with Brfl-targeting siRNA versus control siRNA, plotted and colored as in (A). **D**. MSigDB Hallmark gene-set enrichment for the genes upregulated upon Brfl depletion in lAV-infected cells, plotted as in (B). **E**. Heatmap of log_2_ fold change (Brfl siRNA vs. control siRNA, HSV-1 or IAV infected) for significantly upregulated genes (p-value < 0.05) that overlap one or more MSigDB Hallmark gene sets associated with TNFa signaling via NF-kB. Genes.

Consistent with the phenotype observed during MHV68 infection, preventing Pol III occupancy by Brf1 depletion during HSV-1 infection induced a set of genes significantly enriched for TNF*α* signaling via NF-*κ*B (**Supplemental Fig. 7A**). These 56 genes included innate immune effectors such as IRF1, RELA, NFKBIA, CXCL1, and CXCL2. In IAV-infected cells, the top-ranked Hallmark categories were instead pro-viral cell-cycle and proliferation programs (E2F targets, G2M checkpoint, MYC targets, and mitotic spindle) (**Supplemental Fig. 7B**). This proliferative signature was driven by canonical E2F/cell-cycle effectors and mirrors our observation that the SINE-proximal population of Polr3A engaged promoters during MHV68 infection of proliferating fibroblasts was enriched for E2F target genes (**Fig. 1G**). Filtereing out these cell-cycle Hallmark gene sets from the input gene lists led to TNF*α* signaling via NF-*κ*B emerging as the dominant, significantly enriched pathway in both infections (HSV-1: adjusted *p* = 2.4 × 10^−11^, 31 genes; IAV: adjusted *p* = 0.042, 22 genes; **Fig. 6C-E**). To confirm that this inflammatory de-repression was infection-specific rather than a baseline consequence of Brf1 knockdown, we performed the same cell-cycle-excluded analysis on mock-infected A549 cells, which instead returned enrichment of oxidative phosphorylation, MYC targets, DNA repair, and mTORC1 signaling, with no TNF*α*/NF-*κ*B signal (**Supplemental Fig. 7C**). Together, these data establish that Brf1-dependent restraint of TNF*α*/NF-*κ*B target gene expression is conserved across MHV68, HSV-1, and IAV infections, and across both murine and human cells.

### TBP dependency underlies RNA Polymerase III regulation of inflammatory gene expression

Finally, we searched for feature(s) that might under-lie the regulation of specific protein-coding genes by Pol III. Scanning the promoters of Brf1-restrained genes for canonical core-promoter elements revealed that they were significantly enriched for TATA-box sequences (**Fig. 7A**). This was true across multiple gene groupings: the human inflammatory subset (TNF*α*/NF-*κ*B members, n=48), the human HSV-1 and IAV shared (n=96) and combined (n=1,129) sets, and the MHV68 mouse restrained set (n=373). The enrichment scaled with inflammatory specificity (1.4-fold in the broad set to 5.7-fold in the inflammatory subset, where 23% of promoters carried a TATA-box compared to 5% genome-wide; *p* = 2 × 10^−5^; **Fig. 7A**). Enrichment of TATA-boxes is notable because TATA-binding protein (TBP) is part of both the Pol II initiation factor TFIID complex and the Pol III initiation factor complex TFIIIB, where it is bound directly by Brf1 (Gouge et al., 2017; Hernandez, 1993; Juo et al., 2003; Taggart et al., 1992). The Initiator element, which recruits TFIID via TBP-independent TAF1/TAF2 contacts, was not enriched in any restrained set, indicating that the architectural signature is specific to TBP-binding sequences rather than to core-promoter elements in general.

**Figure 7.**
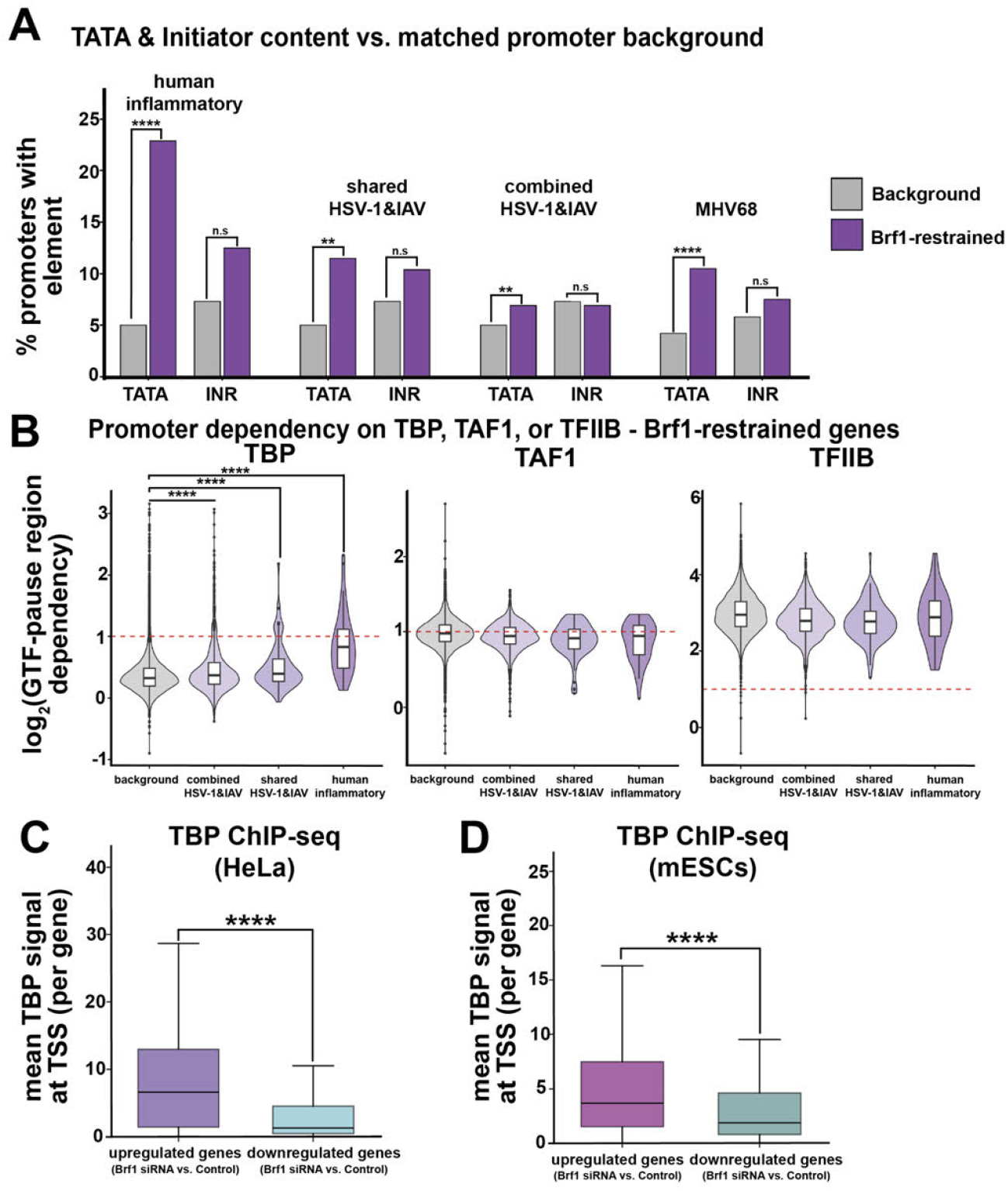
TBP-dependency and occupancy characterize Brfl-restrained inflammatory genes. **A**. Promoter architectural features of Brfl-restrained gene sets. Bar plot showing the percentage of promoters (TSS −50 to +50 bp) containing a canonical TATA-box clement (left) or Initiator (Inr) element (right) in Brfl-restrained gene sets compared to matched-promoter background universes. Gene sets: human inflammatory subset (TNFo/NF-kB Hallmark members of human restrained, n=48), HSV-l&IAV shared restrained (n=96), all human restrained (HSV-1 and IAV upregulated upon Brfl depletion, n= 1,129), and MHV68 mouse restrained (n=373). Promoter matched backgrounds: human = all genes testable in the human DE analysis (n=13,303); MHV68 = all genes testable in the MHV68 DE analysis (n=13,367). Statistical comparison by two-sided Fisher exact test; ^* * * *^ P < IO ^−4, * * *^ P < 10^−3, * *^ P< 10^−2^,^*^ P < 0.05, ns, not significant. **B**. Selective dependence of Brfl-restrained promoters on TBP, but not on TAF1 or TFIIB, in published acute degron-depletion PRO-Seq data (Santana et al., 2022). Violin and box plots show log2-transformed per-gene pause-region dependency (DMSO/dTAGV-1 ratio of PRO-Seq 5’ end reads in the pause region) for the three Pol II general transcription factors TBP, TAF1, and TFIIB, as quantified by Santana et al. 2022 in human HAP1 cells after 2 h dTAGV-1-mediated depletion. Violins depict the full distribution; overlaid boxes show median and interquartile range. Red dashed line, logz(2): Santana et al. TBP-dependent threshold (≤ 2-fold reduction in nascent transcription upon factor depletion). Statistical testing by two-sided Fisher’s exact test on the proportion of TBP-dependent promoters (dep ≥ 2) and one-sided Wilcoxon rank-sum test on continuous dependency values, test set versus human background, excluding the test set. For TBP, inflammatory subset: 34.9% TBP-dependent versus 4.2% in universe (OR = 12.1, Fisher P = 1.6 x IO^−10^; Wilcoxon P = 1.5 x 10^−14^); shared: OR = 3.4, P = 9.4 x 10^−4^; all restrained: OR = 2.6, P= 1.7 x 10^−12^. C. Elevated baseline TBP occupancy at Brfl-restrained promoters in human cells. Per-gene TBPChlP-seq signal at the transcription start sites of expression-matched Brfl-restrained (up) versus Brfl-dependent (down) genes from the combined HSV-1 and IAV restrained set. To control for the dependence of promoter TBP occupancy on transcriptional output, each upregulated gene was matched to a downregulated gene of the nearest baseline (Mock) expression. The TBP signal is from published HeLa S3 ChlP-seq (ENCODE GSM935606), quantified as the mean signal per gene across the TSS ±500 bp window. Boxes show median and interquartile range; whiskers show 1.5 x IQR; outliers omitted for clarity. Two-sided Mann-Whitney U test, p = 4.5 x 10^−9^ **D**. As in (C), using an independent mouse embryonic stem cell TBP ChlP-seq dataset (Kwan et al., 2023), confirming the occupancy difference across species and ChIP datasets. Expression-matched Brfl-restrained (up) versus Brfl-dependent (down) promoters after matching and zero-signal removal; per-gene mean TBP signal across the TSS ± 500 bp window quantified as in (C). Boxes show median and interquartile range; whiskers show 1.5 x IQR; outliers omitted. Two-sided Mann-Whitney U test, p = 5.8 x 10^−7^.

To test whether these promoters functionally depend on TBP, we leveraged published acute degron-mediated depletion PRO-Seq data for TBP, TAF1, and TFIIB in human HAP1 cells (Santana et al., 2022), which quantified per-gene dependency on each factor (**Fig. 7B**). TBP dependency was selectively enriched in a manner that paralleled inflammatory specificity, reaching a 12-fold over-representation of TBP-dependent promoters in the inflammatory subset (34.9% vs. 4.2% in the background set; *p* = 1.6 × 10^−10^; Wilcoxon *p* = 1.5 × 10^−14^). In contrast, TAF1 dependency was non-significantly depleted at all Brf1-restrained sets, and TFIIB dependency was uniformly high across both restrained sets and the background, confirming these are conventional Pol II promoters with no exceptional requirement for the general initiation machinery. Brf1-restrained promoters are thus more TBP-dependent.

We lastly compared TBP occupancy from published human (ENCODE, GSM935606) and mouse (Kwan et al., 2023) ChIP-seq datasets at the TSSs of Brf1-restrained protein-coding genes, matching each gene set to an equivalent baseline (uninfected) expression level so that the difference reflected promoter architecture rather than activity. The Brf1-restrained promoters we identified in HSV-1 and IAV-infected human and MHV68-infected mouse cells showed significantly higher TBP occupancy at the TSS than matched Brf1-dependent promoters (**Fig. 7C-D**). Because both datasets derive from uninfected cells, this reflects an intrinsic property of Brf1-restrained promoters rather than a consequence of infection. Thus, inflammatory genes whose expression is restrained by Pol III during infection are enriched for TATA boxes and TBP occupancy at their promoters, and exhibit TBP-dependent expression, suggesting that competition between Pol II and Pol III for TBP underlies their Pol III-restrained expression signature.

## DISCUSSION

We have identified a pathogen-induced regulatory interplay between Pol II and Pol III at protein-coding gene promoters. In uninfected cells, Pol III occupies TBP-bound, TATA-box-enriched promoters of immune, pro-inflammatory, and stress-responsive genes, keeping the chromatin accessible and likely promoting low-level transcription by Pol II. This could create an environment in which these genes are primed for induction upon pathogen exposure. However, during infection, as Pol II rapidly occupies and induces these loci, Pol III may obstruct or compete with Pol II for shared machinery. Thus, in response to infection, a substantial pool of Pol III moves from the promoters of immune-related genes to SINE retrotransposons. Not all Pol III is moved, however, and the remaining Pol III tempers the degree of Pol II transcription of these loci, perhaps to avoid an over-exuberant inflammatory response. Pol III activity also influences the diversity of isoforms and the processing of mRNAs encoding innate immune and stress response factors. Together, these findings indicate that Pol III occupancy at protein-coding gene promoters represents a context-dependent regulatory process that is actively remodeled during pathogenic stress to promote host defense.

The observation that Pol III is broadly enriched at or near protein-coding gene promoters and supports Pol II gene expression in uninfected cells agrees with previous findings in human cancer cell lines and murine cells (Jiang et al., 2022; K C et al., 2024). Yet, our finding that Pol III occupancy is dynamically regulated in response to infection suggests that it may be particularly important for finely tuned transcriptional responses to cellular stress. Pol III is required to sustain essential housekeeping non-coding RNA production during stressors such as viral infection, nutrient deprivation, heat, or hypoxia; yet it must also respond to changing needs in cellular growth, protein synthesis, and energy expenditure (Ernens et al., 2006; Kulaberoglu et al., 2021; Moir and Willis, 2013; Nguyen et al., 2025; Willis and Moir, 2018). We propose that one way that cells accomplish this is by redistributing existing Pol III pools across distinct genomic loci. This provides a flexible mechanism to couple stress responses with global transcriptional reprogramming without compromising core Pol III functions.

SINEs play increasingly appreciated roles in gene regulation and host defense. Our data here indicate they can serve as reservoirs for Pol III, either as an evolved regulatory feature or because of their abundance and high affinity for the Pol III transcriptional machinery during infection and stress (Fornace and Mitchell, 1986; Jang and Latchman, 1989; Liu et al., 1995; Panning and Smiley, 1993; Panning and Smiley, 1994; Williams et al., 2004). SINE sequences have also been exapted for other aspects of transcriptional regulation, including functioning as enhancers, alternative promoters for Pol II-transcribed genes, boundary elements, and contributors to higher-order chromatin organization (Bourque et al., 2008; Chen et al., 2008; Ferrigno et al., 2001; Horton et al., 2023; Polak and Domany, 2006; Roy-Engel et al., 2005; Su et al., 2014), while Alu elements within mRNAs can induce STAU1-mediated decay (Gong and Maquat, 2011; Gong et al., 2013; Wang et al., 2013). Additionally, SINE-derived non-coding RNAs can activate cytosolic RNA sensors to potentiate innate immune signaling (Aune et al., 2022; Cheng et al., 2020; Guo et al., 2018; Heinrich et al., 2019; Kaneko et al., 2011; Tarallo et al., 2012). Thus, the infection-induced re-localization of Pol III may simultaneously rewire local chromatin landscapes, relieve Pol III-mediated repression at protein-coding gene promoters, and activate immune signaling to promote host defense.

The extensive Brf1-dependent remodeling of mRNA isoforms we observe in both uninfected and infected BMDMs indicates that Pol III activity also affects mRNA processing, including transcription start and end site usage, as well as splicing. Although the underlying mechanisms remain to be determined, we hypothesize that Pol III occupancy can shape the promoter architecture and chromatin environment at Pol II promoters, which are tightly coupled to splicing and RNA stability. Pol III may also compete for or redistribute RNA processing machinery in a context-specific manner. For example, we recently discovered that the mRNA cleavage and polyadenylation machinery can associate with Pol III loci during MHV68 infection, further supporting coordinated regulation of Pol II and Pol III activity and RNA processing (Lari et al., 2025). Furthermore, in mESCs, depletion of the small Pol III subunit Polr3A reduces Pol II occupancy at transcription start sites, decreases Pol II elongation rates, and increases Pol II pausing at a subset of protein-coding genes, changes known to impact mRNA processing (Jiang et al., 2022).

Our finding that Brf1-restrained promoters are architecturally enriched for TATA-boxes, functionally dependent on TBP, and physically marked by elevated baseline TBP occupancy points to a shared requirement for TBP as the molecular basis of Pol III-Pol II crosstalk at these loci. Because TBP is a limiting, shared component of both the Pol II (TFIID) and Pol III (TFIIIB) initiation machineries (Khoo et al., 2014), promoters that depend most heavily on TBP would be the most sensitive to its sequestration by an alternative polymerase. This sequestration is biochemically plausible because Brf1 binds free TBP with higher affinity than TFIIB does in vitro (Schröder et al., 2003), positioning the Pol III machinery to compete favorably for a shared TBP pool. We propose that Pol III occupancy at these promoters draws on this common pool, such that the redistribution of Pol III to SINE retrotransposons during infection releases TBP and lowers a barrier to Pol II initiation, amplifying inflammatory induction. This model links the three features that define the restrained promoter class and supports why TBP-dependence, rather than promoter activity, predicts susceptibility to Brf1-mediated restraint. Such restraint may itself be adaptive, since unchecked TNF*α*/NF-*κ*B and interferon induction drives immunopathology (Liu et al., 2017). By retaining a residual presence at these promoters even as most of the polymerase redistributes to SINEs during viral infection, Pol III may act as a rheostat, permitting a robust response while holding it below a pathological threshold.

We expect the tuning of cellular stress responses by Pol III to be highly context-dependent. Indeed, Pol III binding landscapes vary across cell types and in response to diverse cellular stress conditions (Advani and Ivanov, 2019; Alla and Cairns, 2014). Our integrated analyses support a model in which Pol III promoter occupancy and redistribution are tightly coupled to the cellular response to viral infection. Defining how these dynamics are shaped by distinct stressors and across different cellular contexts will be an important direction for future investigation.

## MATERIALS AND METHODS

### Cell lines

NIH 3T3 (ATCC CRL-1658) and NIH 3T12 (ATCC CCL-164) mouse fibroblast, and human A549 epithelial (ATCC CCL-185) cell lines were maintained in Dulbecco’s modified Eagle’s medium (DMEM; Gibco) with 10% fetal bovine serum (FBS; VWR). BMDMs were differentiated as previously described (Schaller et al., 2020), and experiments involving mice were performed in accordance with a protocol approved by the Institutional Animal Care and Use Committee (IACUC) of the Albert Einstein College of Medicine. Differentiated BMDM cell lines were maintained in BMDM media: Dulbecco’s modified Eagle’s medium (DMEM; Gibco) with 10% fetal bovine serum (FBS; VWR), 1% GlutaMAX Supplement (Gibco), and 62.5 ng/mL recombinant mouse M-CSF (BioLegend). Cell lines were screened regularly for mycoplasma by PCR.

### siRNA nucleofections

Nucleofections of siRNAs into NIH3T3, A549, or BMDM cells were completed as follows: Cells were grown in DMEM with 10% FBS (3T3 and A549) or BMDM media to 90% confluence and were then removed and washed once with Dulbecco’s phosphate-buffered saline (DPBS (Gibco). Nucleofections were done using the Neon Transfection System (Thermo Fisher). 2 × 106 cells were resuspended in 100 *µ*L of buffer R, to which control non-targeting or Brf1 pools of siRNA (ON-TARGETplus SMARTpool siRNA mouse or human - Horizon Discovery) were added to a final concentration of 200nM. This was loaded into a Neon 100-*µ*L pipette tip and a Neon tube with 3 mL of buffer E2, with electroporation parameters set to 1300 V, 20ms, 2 pulses (for NIH3T3), 1200V, 30ms, 2 pulses (for A549), or 1,680 V, 20 ms, 1 pulse (for BMDM). Following electroporation, cells were plated in 10 mL of DMEM with 10% FBS or BMDM media in 10 cm TC-treated plates and incubated at 37°C for 24 hours. siRNA nucleofections were repeated after 24 hours for BMDMs, and cells were plated and incubated at 37°C for another 24 hours.

### Virus preparations and infections

### MHV68 infections

MHV68 was amplified in NIH 3T12 fibroblast cells, and the viral 50% tissue culture infective dose (TCID50) was measured on NIH 3T3 fibroblasts by limiting dilution. NIH 3T3 fibroblasts were infected at the indicated multiplicity of infection (MOI) by adding the required volume of the virus to cells in 5 mL of serum-free DMEM in 10-cm TC-treated plates. BMDMs were infected with the minimal volume of MHV68 required to achieve maximum infection (20% of cells), as determined by titration experiments with GFP-marked MHV68 followed by flow cytometry. For infections in BMDMs, the virus was added to cells in serum-free DMEM for 4 hours. Virus-containing media were then aspirated and replaced with BMDM media.

#### HSV-1 infections

Human herpesvirus 1 strain KOS (HSV-1 KOS; ATCC VR-1493) was propagated in Vero cells, and the viral titer was determined by plaque assay. Cells were infected with HSV-1 at an MOI of 0.1 using a virus volume calculated from plaque assay-determined titers. For infection, the virus was diluted in PBS and incubated with A549 cells for 1 hour. Following adsorption, cells were washed once with PBS, and fresh culture medium was added. Cells were harvested 24 hours post-infection.

#### Influenza A virus infections

Influenza A virus strain A/Puerto Rico/8/1934 (H1N1) (PR8; ATCC VR-1469) was propagated in MDCK cells, and viral titers were determined by the manufacturer using the CEID50 assay. A549 cells were washed twice with PBS and infected with PR8 at an MOI of 1.0 in a minimal volume of PBS for 1 hour at 37°C with gentle rocking every 15 minutes. Following adsorption, the inoculum was removed, cells were washed twice with PBS, and fresh serum-free medium supplemented with 0.3% BSA and 0.5 *µ*g/mL TPCK-treated trypsin was added. Cells were incubated for 24 hours at 37°C prior to harvest.

### Western blotting

To prepare whole-cell lysates for evaluating protein expression of Brf1, cells were washed with cold DPBS (Gibco) followed by lysis with radioimmunoprecipitation assay (RIPA) lysis buffer (50 mM Tris HCl, 150 mM NaCl, 1.0% [vol/vol] NP-40, 0.5% [wt/vol] sodium deoxycholate, 1.0 mM EDTA, and 0.1% [wt/vol] SDS, Roche cOmplete Mini EDTA-free protease inhibitor cocktail). Cell lysates were vortexed briefly, rotated at 4°C for 15 min, and then clarified by centrifugation at 21,000 × g in a tabletop centrifuge at 4°C for 10 min to remove debris. 30 *µ*g of whole-cell lysate were resolved on 4% to 15% mini-PROTEAN TGX gels (Bio-Rad). Transfers to polyvinylidene difluoride (PVDF) membranes (Bio-Rad) were done with the Trans-Blot Turbo transfer system (Bio-Rad). Blots were incubated in 5% milk in TBS with 0.1% Tween 20 (TBS-T) to block, followed by incubation with primary antibodies against anti-Brf1 (Bethyl Laboratories A301-228A, 1;1000), anti-Acod1 (Abcam ab222411, 1;1000), anti-Serpine1 (Invitrogen MA5-17171, 1;1000), anti-Polr3D (1;1000), anti-Polr3I (1:1000), anti-Polr3G (1;1000), anti-Polr3F (1;1000), anti-GAPDH (Invitrogen AM4300, 1;500), and anti-Vinculin antibody (ab-cam ab91459, 1;5,000). Washes were carried out with TBS-T. Blots were then incubated with HRP-conjugated secondary antibodies (Southern Biotechnology, 1:5,000). Washed blots were incubated with Clarity Western ECL substrate (Bio-Rad) for 5 min and visualized with a ChemiDoc imager (Bio-Rad).

### ChIP-seq visualization and promoter occupancy analysis

#### Processing

For visualization of Polr3A and TBP ChIP-seq datasets, aligned BAM files mapped to the appropriate reference genome (mm10 for murine datasets and hg38 for human datasets) were converted to Big-Wig files using the bamCoverage function in deep-Tools (Ramírez et al., 2016). For spike-in-normalized datasets, BigWig files were generated using the parameters –binSize 10, –smoothLength 30, –extendReads, – centerReads, and –normalizeUsing None, together with a dataset-specific –scaleFactor corresponding to the reciprocal of the spike-in normalization factor calculated during differential binding analysis. For datasets lacking spike-in controls, the signal was normalized using the sequencing-depth normalization method. Replicate-normalized BigWig files from two biological replicates per condition, unless otherwise indicated, were averaged using bigwigAverage with a 10-bp bin size (-bs 10) to generate a representative signal track for downstream visualization and quantification. Differential ChIP-seq occupancy was analyzed using DiffBind with the DESeq2 statistical framework (Love et al., 2014; Stark, 2011). Each ChIP-seq peak was assigned to a single genomic category by combining gene-structure annotations generated with ChIPseeker (Yu et al., 2015) and a transcription start site region of −1,000 to +1,000 bp, and intersecting with RepeatMasker repeat annotations using BEDTools (Smit, 2013-2015). Peaks overlapping both gene-associated and repetitive genomic features were assigned according to the predefined annotation hierarchy used for downstream analyses.

#### Heatmaps & metagene profiles

Heatmaps and metagene profiles were generated from replicate-averaged BigWig files using deepTools computeMatrix reference-point. Analyses were centered on annotated transcription start sites (TSSs) of protein-coding genes using the corresponding genome annotation. Blacklisted genomic regions were excluded when available, and matrices were generated using the options –skipZeros and – missingDataAsZero. ChIP-seq signal was plotted as histograms with 10-bp bins across the indicated genomic intervals surrounding the TSS. Heatmaps were ranked by average ChIP-seq occupancy unless otherwise indicated. Metagene analyses of differentially expressed gene sets were performed using the same replicate-averaged Big-Wig files and visualization parameters.

#### Visualization

Replicate-averaged, normalized Big-Wig files were displayed in the Integrative Genomics Viewer (IGV) (Robinson et al., 2011). Representative loci were selected from differentially bound Polr3A regions, inflammatory gene promoters, interferon-stimulated genes, SINE-associated loci, and representative TBP-bound promoters. For comparisons between conditions, tracks were visualized using identical y-axis scales within each locus.

#### Promoter occupancy quantification

To quantify promoter-associated Polr3A or TBP occupancy, the mean ChIP-seq signal from replicate-averaged BigWig files was calculated within a ± 500 bp window surrounding each annotated protein-coding gene TSS. Promoters were ranked by occupancy and stratified into quintiles for downstream analyses. These promoter occupancy scores were used to compare chromatin accessibility changes measured by ATAC-seq and to assess relationships between promoter occupancy, promoter architecture, and transcriptional responses to Brf1 depletion or viral infection.

#### Total chromatin occupancy analyses

For calculations of total Polr3A chromatin occupancy, signal intensity was quantified across the union set of Polr3A peaks identified by MACS2 (Zhang et al., 2008). Integrated signal was calculated as the mean normalized ChIP-seq signal multiplied by peak length and summed across all peaks within each condition. Peak-level comparisons between conditions were performed using replicate-averaged spike-in-normalized BigWig files.

Publicly available ChIP-seq datasets were obtained from NCBI’s Gene Expression Omnibus and are accessible under the GEO accession numbers here: Polr3A (**GSE288677**) (Lari et al., 2025), and TBP (human – **ENCODE GSM935606**, mouse - **GSE172401**) (Kwan et al., 2023). The publicly available Brf1 and Brf2 ChIP-seq datasets were obtained from the ArrayExpress database under accession **E-MTAB-767** (Carrière et al., 2011).

### RNA-sequencing

#### RNA extraction

Total RNA was extracted from cells in triplicate using TRIzol reagent (Invitrogen), and total RNA integrity was assessed by Agilent TapeStation using RNA ScreenTape. Matched RNA from murine primary BMDMs was used for both short-read and long-read sequencing; RNA from human A549 cells was used for 3’ mRNA-seq.

#### Short-read RNA-seq library preparation (murine primary BMDMs)

For each sample, 100 ng of total RNA was spiked with ERCC RNA Spike-In Mix 1 (Thermo Fisher Scientific), and libraries were prepared using the KAPA RNA HyperPrep Kit with RiboErase (HMR) (Roche) with KAPA Unique Dual-Indexed adapters, according to the manufacturer’s instructions. Pooled libraries were sequenced on an Illumina NovaSeq X (150 bp, paired end).

#### Short-read read processing and differential expression

RNA-seq reads were processed using HT-Stream (v1.4.1) to remove adapter sequences and low-quality reads before alignment to the mouse (mm10) reference genome using STAR (v2.7.11b). Aligned reads were filtered with samtools (v1.22.1) to retain only properly paired, uniquely mapped, non-supplementary reads. Gene-level read counts were generated with featureCounts (v2.1.1) by quantifying reads overlapping annotated exons. ERCC RNA spike-in controls were used for normalization to account for technical variation in library preparation and sequencing depth. Differential gene expression analysis was subsequently performed using the limma-voom pipeline with ERCC-derived normalization factors. Genome-wide RNA-seq coverage tracks were generated from filtered alignment files in bigWig format using bamCoverage from deepTools, with signal scaled using the corresponding ERCC-derived normalization factors. Replicate-level bigWig files were averaged within each experimental condition using bigwigCompare from deepTools to generate condition-level tracks for genome-browser visualization.

### Long-read RNA-seq library preparation (murine primary BMDMs)

Full-length cDNA libraries were generated from the same total RNA used for short-read sequencing using the PacBio SMRTbell Express Template Prep Kit 2.0 (100-938-900). Briefly, 300ng of total RNA was reverse-transcribed and amplified to generate full-length cDNA, from which SMRTbell libraries were constructed. Libraries were sequenced on a PacBio Sequel II instrument.

### Long-read read processing and isoform analysis

PacBio Iso-Seq data were processed using the IsoSeq3 suite (v3.4.0) following the cDNA_Cupcake Iso-Seq bioinformatics workflow (https://github.com/Magdoll/cDNA_Cupcake/).

Primer sequences were removed from circular consensus sequencing (CCS) reads using lima, followed by removal of poly(A) tails and concatemers using IsoSeq3 refine to generate full-length non-concatemer (FLNC) reads. FLNC reads were clustered using IsoSeq3 cluster to generate high-confidence transcript isoforms, which were aligned to the mouse mm39 reference genome using pbmm2. Aligned isoforms were classified and annotated at the gene, transcript, and splice-junction levels using SQANTI3 (v4.1) (Tardaguila et al., 2018). Isoform-level read counts were generated by mapping FLNC read identifiers to their sample of origin and corresponding SQANTI3-classified transcript identifiers. Differential isoform expression analysis was performed in R using the limma-voom pipeline (v3.52.1) (Law et al., 2014; Ritchie et al., 2015). Count data were normalized using the trimmed mean of M-values (TMM) method (Robinson and Oshlack, 2010), lowly expressed transcripts were filtered prior to statistical testing, and P values were adjusted for multiple comparisons using the Benjamini–Hochberg procedure (Benjamini and Hochberg, 1995).

#### 3’ mRNA-seq library preparation and sequencing (human A549)

Total RNA was extracted from A549 cells in triplicate and spiked with ERCC RNA Spike-In Mix 1 (Thermo Fisher Scientific) at equal volume. 3’ mRNA-seq libraries were prepared by Plas-midsaurus using a 3’ end RNA-seq workflow and sequenced on Illumina NovaSeq Plus instrument as single-end, stranded reads (90 bp).

#### 3’ mRNA-seq read processing and differential expression

Reads were demultiplexed with BCL Convert (v4.3.6) and fqtk (v0.3.1), trimmed with fastp (v0.24.0; poly-X tail trimming, 3’ quality-based trimming, minimum Phred score 15, minimum length 50 bp), and aligned to GRCh38 (Ensembl release 114) with STAR (v2.7.11b) with non-canonical splice junction removal. BAM files were coordinate-sorted with samtools (v1.21), and PCR/optical duplicates were removed using UMI-based deduplication with UMICollapse (v1.1.0). Gene-level 3’ counts were assigned using featureCounts (Subread v2.1.1; exons and 3’ UTR features, strand-specific, fractional multi-mapping assignment, grouped by gene_id). Approximately 10 million deduplicated reads were obtained per sample. Differential expression between control and Brf1-depleted samples during HSV-1 or influenza A virus infection was determined using DESeq2 (v1.48.2) with ERCC spike-in normalization, applying thresholds of (padj < 0.05). ERCC size factors were computed as the ratio of observed to expected spike-in counts per sample and used to assess infection-associated global expression changes; ERCC normalization was not applied to Brf1siRNA versus control siRNA contrasts due to systematic variation in spike-in recovery between siRNA conditions.

#### RNA-seq visualization

Volcano plots/heatmaps were generated in R (v4.5.1) using ggplot2 and ComplexHeatmap from the differential expression results above. Gene set enrichment (GSEA) was performed as described below.

### Promoter architecture analysis

Promoter sequences of Brf1-restrained and control gene sets were defined as regions surrounding the annotated TSS in the mm10 or hg38 reference genomes. TATA-box enrichment was assessed by PWM scanning using the Bucher 1990 (Ambrosini et al., 2018) TATA matrix (min.score = 75% of maximum) within the −40 to −20 bp region relative to the TSS, and Initiator (Inr) motif enrichment was assessed using the YYANWYY consensus PWM (min.score = 85%) within the ^−^ 2 to +5 bp region (Ambrosini et al., 2018; Javahery et al., 1994). Brf1-restrained gene sets were compared with species-matched background sets consisting of all genes testable in the corresponding differential expression analysis. Statistical significance of promoter-feature enrichment was evaluated using two-sided Fisher’s exact tests.

### Gene ontology and MSigDB Hallmark analysis

#### Over-representation analysis (ORA)

of differentially regulated gene sets was performed using the clusterProfiler R package (v4.16; Yu et al., OMICS 2012). For each comparison, query gene lists (e.g., genes up- or downregulated in siBrf1 vs siCtrl; genes with lost or gained Pol III occupancy) were tested against a background universe consisting of all genes detected in the corresponding RNA-seq experiment (mean count > 0 across all samples) or, for ChIP-seq-based gene sets, all GENCODE vM25 protein-coding genes whose promoters were assayed.

#### MSigDB Hallmark enrichment

The mouse-native MSigDB Hallmark collection (50 gene sets) was retrieved using the msigdbr R package (v25; collection = “MH”, db_species = “MM”), which returns the Hallmark gene sets pre-converted to mouse orthologs and using NCBI Entrez gene IDs (column ncbi_gene). Hallmark enrichment was computed with clusterProfiler::enricher using the hypergeometric test, with the following parameters: pvalueCutoff = 0.05, qvalueCutoff = 0.2, pAdjustMethod = “BH”, minGSSize = 10, maxGSSize = 500. Reported P-values are Benjamini–Hochberg-adjusted. Gene Ontology Biological Process enrichment. GO BP over-representation was computed using clusterProfiler::enrichGO with OrgDb = org.Mm.eg.db (v3.22), ont = “BP”, pvalueCutoff = 0.05, qvalueCutoff = 0.2, pAdjustMethod = “BH”, minGSSize = 10, maxGSSize = 500, and gene identifiers in ENTREZID format. Redundant GO terms were collapsed using clusterProfiler::simplify with cutoff = 0.7 and measure = “Wang” to remove semantically similar terms. When the unfiltered enrichment output exceeded 200 terms, results were pre-filtered to retain only terms with adjusted P < 0.01 before simplification to avoid memory issues during semantic similarity computation. Isoform-level enrichment. For the long-read Iso-Seq dataset, differentially expressed isoforms (Brf1siRNA vs Control siRNA, separately in mock and MHV68 conditions) were first collapsed to unique gene symbols, and ORA was performed as above using the gene-collapsed lists.

#### Visualization

Dotplots were generated using enrichplot::dotplot (v1.30) showing the top 10 (Hallmark) or top 15 (GO BP, post-simplification) categories ranked by adjusted P-value. Dot color encodes adjusted P-value (continuous gradient); dot size encodes the number of query genes in the term (“Count”). Plots were assembled into multi-panel layouts using patchwork (v1.3).

### ATAC-sequencing and data processing

#### Library preparation

ATAC-seq libraries were prepared from NIH 3T3 cells transfected with control or Brf1-targeting siRNA and either mock-infected or MHV68-infected at 24 hours post-infection, using the Zymo-Seq ATAC Library Kit (Zymo Research, D5458) according to the manufacturer’s instructions. Drosophila melanogaster S2 nuclei (Active Motif, 53154) were spiked in at a fixed ratio prior to Tn5 tagmentation to enable spike-in normalization across conditions. Libraries were sequenced as 150 bp paired-end reads on a NovaSeq X instrument at the QB3 Genomics facility (University of California, Berkeley), in biological triplicate.

#### Data processing

Raw read quality was assessed using FastQC (v0.12) and MultiQC (v1.33). Adapter sequences were trimmed using Trim Galore (v0.6.10) with default paired-end parameters. Trimmed reads were aligned to a combined mm10/dm6 reference genome using Bowtie2 (v2.5.5; –very-sensitive –no-mixed –no-discordant -X 2000), with all Drosophila chromosomes prefixed with “dm6_” to distinguish spike-in from mouse reads. Mock-infected samples aligned at 96–98% and MHV68-infected samples at 29–43%, consistent with ~ 60% of infected-sample reads deriving from viral genomic DNA. Read groups were added and PCR duplicates removed using Picard (v3.4.0); duplication rates were 11-27% in MHV68-infected and 17-49% in mock-infected samples. Aligned reads were filtered to retain properly paired reads (SAMtools v1.22.1; -f 0×2 -F 0×4 - F 0×8) with MAPQ≥ 30, excluding mitochondrial DNA, Drosophila spike-in reads, and ENCODE mm10 black-listed regions (BEDTools v2.31.1). Final filtered read counts ranged from 38.9 to 150.1 million read pairs per sample. Accessible chromatin peaks were called per sample using MACS3 (v3.0.4; -f BAMPE -g mm –nomodel –nolambda -q 0.05), yielding 75,080–144,807 peaks per sample. A consensus peakset of 203,862 non-overlapping peaks was generated by merging all twelve narrowPeak files using BEDTools merge. Fragment counts within consensus peaks were quantified across all samples using featureCounts (Subread v2.1.1; -p –countReadPairs -B -C), and peaks with fewer than 10 reads in at least 3 samples were removed, retaining 200,910 peaks. Differential chromatin accessibility analysis was performed in R (v4.5.1) using DESeq2 (v1.48) with a 2 × 2 factorial design (~ siRNA + virus + siRNA:virus). Three pair-wise comparisons were extracted: Brf1 siRNA vs. control siRNA in mock conditions; Brf1 siRNA vs. control siRNA during MHV68 infection (using the combined interaction coefficient); and MHV68 vs. mock in control siRNA cells. Significance was defined as a Benjamini-Hochberg adjusted p-value < 0.05. Variance-stabilized counts (VST) were used for principal component analysis of the top 500 most variable peaks. Peaks were annotated to genomic features using ChIPseeker (v1.44.0) with mm10 UCSC KnownGene annotation and a TSS window of − 3,000 to +3,000 bp. Overlaps between consensus peaks and tRNA genes (UCSC tRNAs.txt) or B1/B2 SINE elements (RepeatMasker mm10 rmsk.txt; B1 elements annotated as “Alu” in mouse RepeatMasker, reflecting their shared evolutionary origin with human Alu elements) were identified using BEDTools intersect. Differential accessibility at tRNA-overlapping and SINE-overlapping peaks was assessed within the DESeq2 framework, cross-referencing peak identifiers with SINE annotations to determine the number and direction of significantly changed peaks in each annotation class.

#### Visualization

Spike-in normalization scaling factors were calculated as 1,000,000 divided by the per-sample Drosophila dm6 read count (scaling factors ranged from 0.131 to 0.825). Spike-in normalized and CPM-normalized bigWig files were generated using deepTools bamCoverage (v3.5.6; –binSize 10 –extendReads). Given that viral DNA competed with Drosophila spike-in reads for sequencing depth in MHV68-infected samples, spike-in normalization was used for within-condition comparisons (mock vs. mock, MHV68 vs. MHV68). Condition-averaged bigWig files were generated using deepTools bigwigAverage across the three biological replicates per condition. Genome-wide heatmaps of ATAC-seq log_2_(Brf1 siRNA/control siRNA) at protein-coding gene TSSs (± 3 kb) were generated using deepTools computeMatrix (reference-point –referencePoint TSS -b 3000 -a 3000 –binSize 10) and plotHeatmap, centered on 54,721 protein-coding gene TSS coordinates from GENCODE vM25 annotation (gencode.vM25.annotation.gtf, transcript_type “protein_coding”), with rows sorted by mean log_2_FC signal. ATAC-seq log_2_(Brf1 siRNA/control siRNA) at individual protein-coding gene promoters (TSS ± 500 bp) was stratified by Polr3A ChIP-seq occupancy quintile or by differential expression status, and statistical comparisons were performed using two-sided Mann-Whitney U tests. Absolute mean ATAC signal per Polr3A occupancy quintile was computed from spike-in-normalized bigWig ± 500 bp windows. The number of significantly differentially accessible peaks (adjusted p < 0.05) overlapping tRNA genes or B1/B2 SINE elements was quantified across three pair-wise DESeq2 comparisons and visualized as bar plots using ggplot2 in R. All ATAC-seq analyses are from n=3 biological replicates per condition; Polr3A ChIP-seq data used for comparison are from n=2 biological replicates. Genome browser tracks were visualized using the Integrative Genomics Viewer (IGV, v2.16).

### Analysis of TBP, TAF1, and TFIIB dependency from published degron PRO-seq data

To assess the dependence of Brf1-restrained promoters on individual Pol II general transcription factors, we re-analyzed published precision run-on sequencing (PRO-seq) data from human HAP1 cells engineered for dTAG-mediated acute degradation of TBP, TAF1, or TFIIB (Santana et al., 2022; publicly available at GEO accession **GSE194153**). For each factor, we used the per-gene pause-region quantifications generated by the authors using truQuant, which counts PRO-seq 5’-end reads in the promoter-proximal pause region. Per-gene factor dependency was defined as the ratio of pause-region signal in vehicle-treated (DMSO) versus depleted (dTAGV-1, 2 h) conditions, with higher values indicating a greater reduction in nascent transcription upon factor depletion. Following Santana et al., a gene was classified as factor-dependent if its DMSO/dTAGV-1 ratio was ≥ 2 (log_2_ dependency ≥ 1). Brf1-restrained gene sets, the human inflammatory subset (TNF*α*/NF-*κ*B members, n = 48), the HSV-1 and IAV shared restrained set (n = 97), and the combined human restrained set (n = 1,143), were mapped to the truQuant gene list by gene symbol (inflammatory, 43 of 48 mapped; shared, 83 of 97; combined restrained, 920 of 1,143). All genes quantified in the truQuant list and testable in the human differential-expression analysis served as the background universe (9,100 of 13,303 mapped). For each factor and gene set, enrichment was assessed two ways: a two-sided Fisher exact test on the proportion of factor-dependent promoters (dependency ≥ 2) in the test set versus the background universe (excluding test-set genes), and a one-sided Wilcoxon rank-sum test on the continuous per-gene dependency values (test set versus background, excluding test-set genes). Analyses were performed identically for TBP, TAF1, and TFIIB.

### Data, Materials, and Software Availability

The sequencing datasets generated in this study have been submitted to the NCBI Gene Expression Omnibus (GEO) and are currently under accession processing. The GEO accession number will be added to the manuscript once available and prior to journal publication.

## Acknowledgements

We thank all current and past members of the Glaunsinger and Coscoy Labs. We thank Leah Gulyas and Devon Jeltema for their careful reading of the manuscript. We thank Bradley Jenner at the UC Davis Genome Center Bioinformatics Core for support with long-read RNA-seq data analyses. Sequencing was performed through the Vincent J. Coates Genomics Sequencing Laboratory at UC Berkeley (QB3 Genomics, UC Berkeley, Berkeley, CA, RRID:SCR_022170). This work is funded by National Institutes of Health (NIH) R01CA136367 and R01AI122528 to B.A.G., who is also an investigator of the Howard Hughes Medical Institute.

## Author Contributions

A.L. and B.A.G. designed research; A.L., S.B.S., S.B., and M.N. performed research; A.L., S.B.S., and B.A.G. analyzed data; A.L. and B.A.G. wrote the paper.

## Supplemental Information

Supplemental Figures 1-7

Supplemental Tables 1-6

