## Supplemental Figures for "Promoter-associated RNA polymerase III shapes RNA polymerase II-dependent inflammatory gene expression during viral infection"

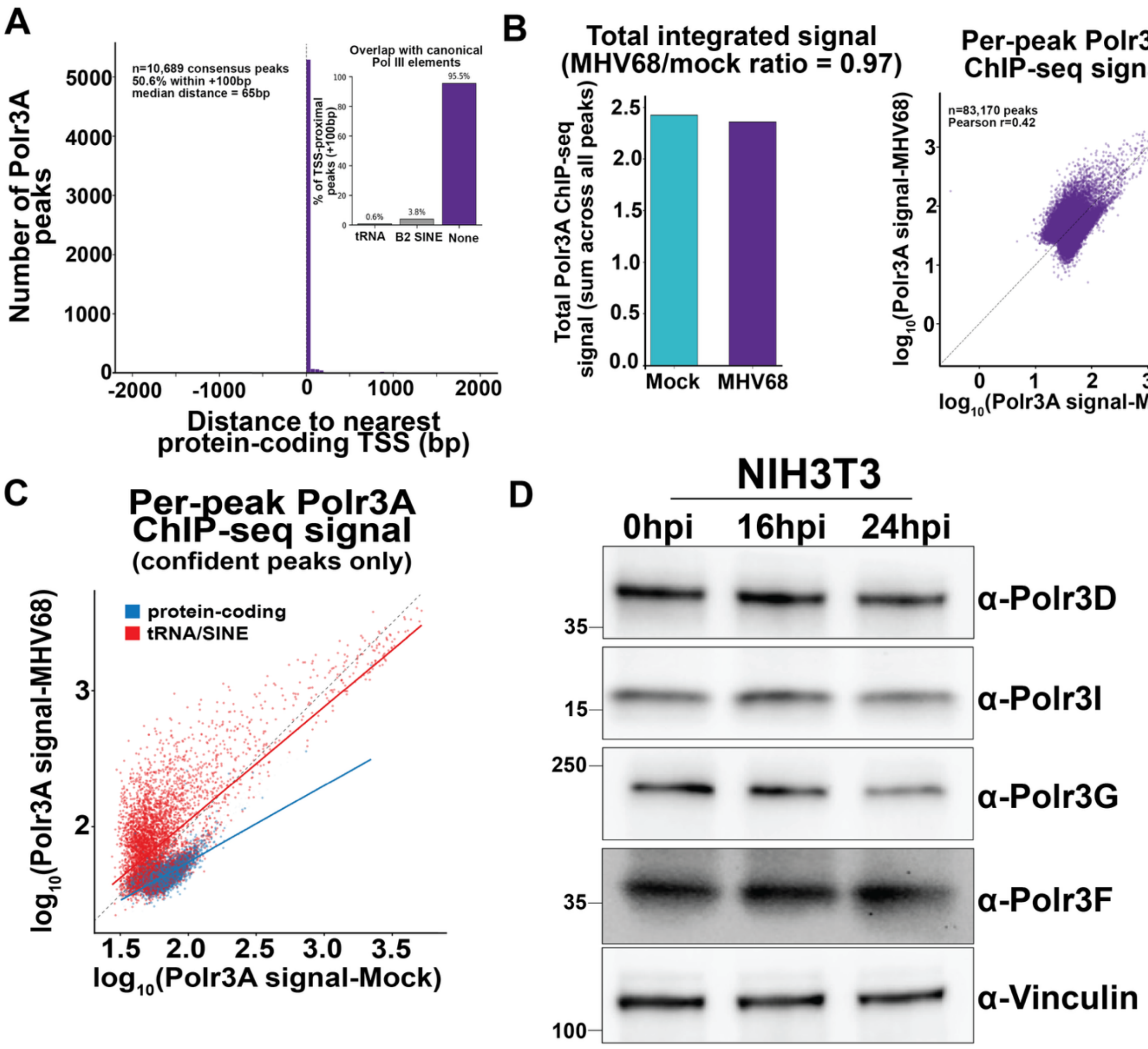


**Supplemental Figure 1. RNA polymerase III ChIP-seq signal and subunit protein abundance do not change during MHV68 infection. A.** Distribution of mock-condition Polr3A consensus peaks (n = 10,689; intersection of two biological replicate MACS2 (Zhang et al., 2008) peak calls) by signed distance to the nearest protein-coding gene TSS. Main plot: ±2 kb zoom histogram with dashed line at the TSS. Annotation: total peak count, % within ±100 bp, and median absolute distance. Inset: bar plot of repeat-element overlap among TSS-proximal (±100 bp) peaks (n = 5,406), showing the percentages overlapping annotated tRNA genes (GtRNAdb) (Chan and Lowe, 2016), B2 SINE retrotransposons (RepeatMasker B2_Mm subfamilies (Smit, 2013-2015)), or neither category. **B.** Conservation of total Polr3A ChIP-seq signal during MHV68 infection. Left: bar plot of total integrated Polr3A ChIP-seq signal across all peaks for each condition (sum of mean signal × peak length). MHV68/Mock ratio = 0.97. The ChIP-seq signal is from spike-in-normalized BigWig files. Right: scatter plot of $\log_{10}$-transformed Polr3A signal per peak in mock (x-axis) vs MHV68-infected (y-axis) NIH3T3 fibroblasts. Each point represents one peak from mock and MHV68 Polr3A MACS2 (Zhang et al., 2008) peak sets (n = 83,170). Dashed line, y = x. Pearson correlation is computed on $\log_{10}$-transformed signal values. **C.** Scatter plot of $\log_{10}$-transformed Polr3A signal per peak in mock (x-axis) vs. MHV68-infected (y-axis) NIH3T3 fibroblasts, colored by peak category (tRNA/SINE, red; protein-coding gene promoter, blue; other, grey; see Methods). Each point represents one peak from the mock and MHV68 Polr3A MACS2 (Zhang et al., 2008) peak sets (n = 83,170). Dashed line, y = x. Colored lines show linear fits restricted to peaks independently detected as significant in both mock and MHV68 (tRNA/SINE, n = 5,468, slope = 0.84, 95% CI 0.82–0.86; protein-coding, n = 3,311, slope = 0.56, 95% CI 0.54–0.58). **D.** NIH3T3 cells were mock-treated or infected with MHV68 at an MOI of 5. At the indicated time points, cells were harvested and lysed to extract total protein, which was then analyzed by Western blotting using antibodies against Polr3D, Polr3I, Polr3G, Polr3F, and Vinculin (loading control).

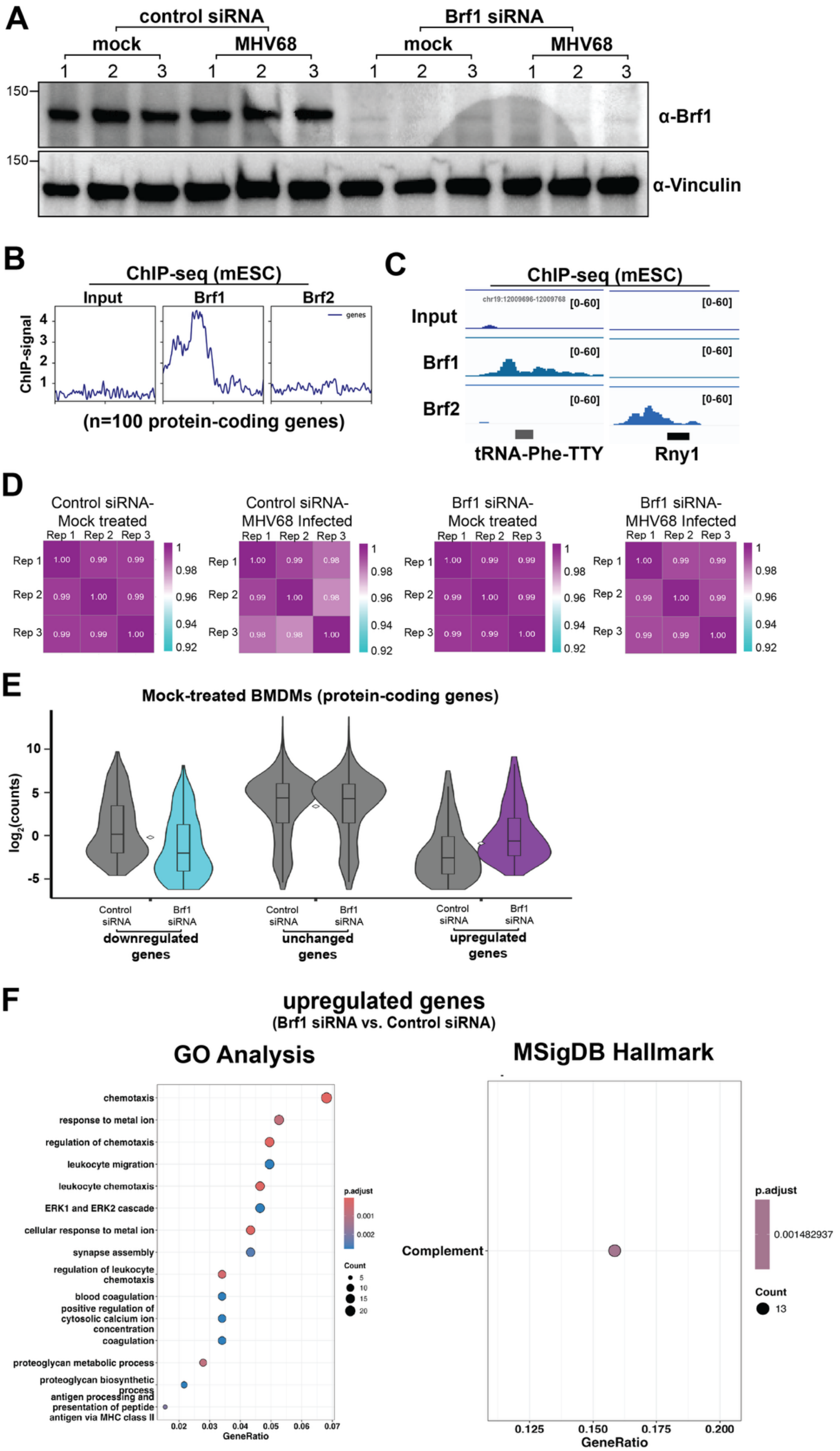
A
control siRNA
Brf1 siRNA
mock
MHV68
mock
MHV68
1 2 3 1 2 3 1 2 3 1 2 3
150
α-Brf1
150
α-Vinculin
B
ChIP-seq (mESC)
Input
Brf1
Brf2
genes
ChIP-signal
(n=100 protein-coding genes)
C
ChIP-seq (mESC)
Input
Brf1
Brf2
[0-60]
tRNA-Phe-TTY
Rny1
D
Control siRNA-Mock treated
Control siRNA-MHV68 Infected
Brf1 siRNA-Mock treated
Brf1 siRNA-MHV68 Infected
Rep 1
Rep 2
Rep 3
E
Mock-treated BMDMs (protein-coding genes)
log2(counts)
Control siRNA
Brf1 siRNA
downregulated genes
unchanged genes
upregulated genes
F
upregulated genes
(Brf1 siRNA vs. Control siRNA)
GO Analysis
chemotaxis
response to metal ion
regulation of chemotaxis
leukocyte migration
leukocyte chemotaxis
ERK1 and ERK2 cascade
cellular response to metal ion
synapse assembly
regulation of leukocyte chemotaxis
blood coagulation
positive regulation of cytosolic calcium ion concentration
coagulation
proteoglycan metabolic process
proteoglycan biosynthetic process
antigen processing and presentation of peptide antigen via MHC class II
GeneRatio
p.adjust
Count
MSigDB Hallmark
Complement
GeneRatio
p.adjust
0.001482937
Count
13

**Supplemental Figure 2. Brf1 knockdown in primary BMDMs alters protein-coding gene expression. A.** Primary BMDMs transfected with control non-targeting or Brf1 targeting siRNAs were mock-treated or infected with MHV68 at an MOI of 5 in triplicate. At 48 hpi, cells were harvested and lysed to extract total protein, which was then analyzed by Western blotting using antibodies against Brf1 and Vinculin (loading control). **B.** Metagene plot displaying Brf1 or Brf2 ChIP-seq signal (Nguyen et al., 2025) across protein-coding genes. Protein-coding genes were selected based on the top 100 genes with the most Pol III ChIP-seq signal from Fig. 1. ChIP-seq signal was plotted as a histogram with 10 bp bins from -1 to +1kbp around the transcription start site (TSS) with 20 bins/gene. Heatmaps are ranked by ChIP-seq coverage values. **C.** Brf1 or Brf2 ChIP-seq (Nguyen et al., 2025) coverage across a select tRNA (type II promoter) or Y RNA gene (type III promoter). Alignment files were visualized in the Integrative Genome Viewer (IGV) (Robinson et al., 2011). Genomic coordinates and Y-axis maximum and minimum values are within brackets. **D.** Spearman correlation coefficients between biological replicates from RNA-seq experiment in primary BMDMs transfected with control non-targeting or Brf1 targeting siRNAs that were mock-treated or infected with MHV68 at an MOI of 5 for 48 hours in triplicate **E.** Violin box plots showing spike-in normalized average expression for genes classified by their differential expression status from (Figure 2B - downregulated, unchanged, upregulated) and stratified by conditions (control siRNA or Brf1 siRNA). Violins depict the full distribution; overlaid boxes show the median and interquartile range; and white diamonds mark the mean. **F.** Gene Ontology Biological Process (GO:BP) (Yu et al., 2012) enrichment (left) and MSigDB Hallmark (Liberzon et al., 2015) gene-set enrichment (right) of Brf1-dependent upregulated protein-coding genes in BMDMs. Dot size indicates gene count, and dot color indicates the Benjamini-Hochberg (BH)-adjusted p-value. GO enrichment simplified by semantic similarity (clusterProfiler::simplify, cutoff = 0.7) to remove redundant terms. Top 15 terms shown per direction. Enrichment was performed with clusterProfiler::enricher using the mouse-native Hallmark collection (msigdbr collection “MH”).

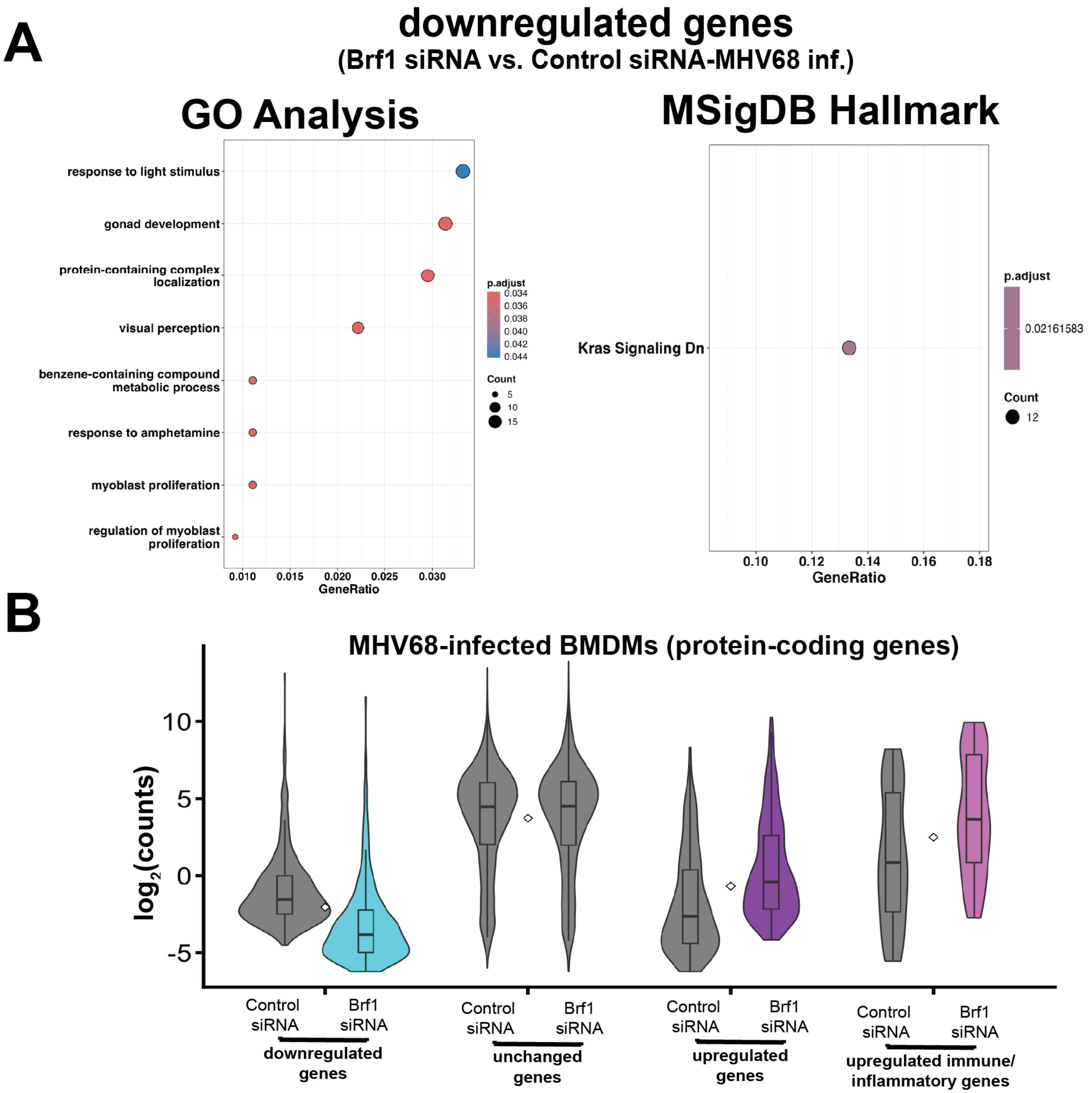


**Supplemental Figure 3. Brf1 knockdown in MHV68-infected primary BMDMs results in upregulation of pro-inflammatory genes. A.** Gene Ontology Biological Process (GO:BP) (Yu et al., 2012) enrichment and MSigDB Hallmark (Liberzon et al., 2015) gene-set enrichment of Brf1-dependent downregulated protein-coding genes in MHV68-infected BMDMs. GO enrichment simplified by semantic similarity (clusterProfiler::simplify, cutoff = 0.7) to remove redundant terms. Top 15 terms shown per direction. Dot size indicates gene count, and dot color indicates the Benjamini-Hochberg (BH)-adjusted p-value. Enrichment was performed with clusterProfiler::enricher using the mouse-native Hallmark collection (msigdbr collection "MH"). **B.** Violin box plots showing spike-in normalized average expression for genes classified by their differential expression status from Figure 3A (downregulated, unchanged, upregulated, or upregulated immune/inflammatory genes and stratified by conditions (control siRNA or Brf1 siRNA). Violins depict the full distribution; overlaid boxes show the median and interquartile range; and white diamonds mark the mean.

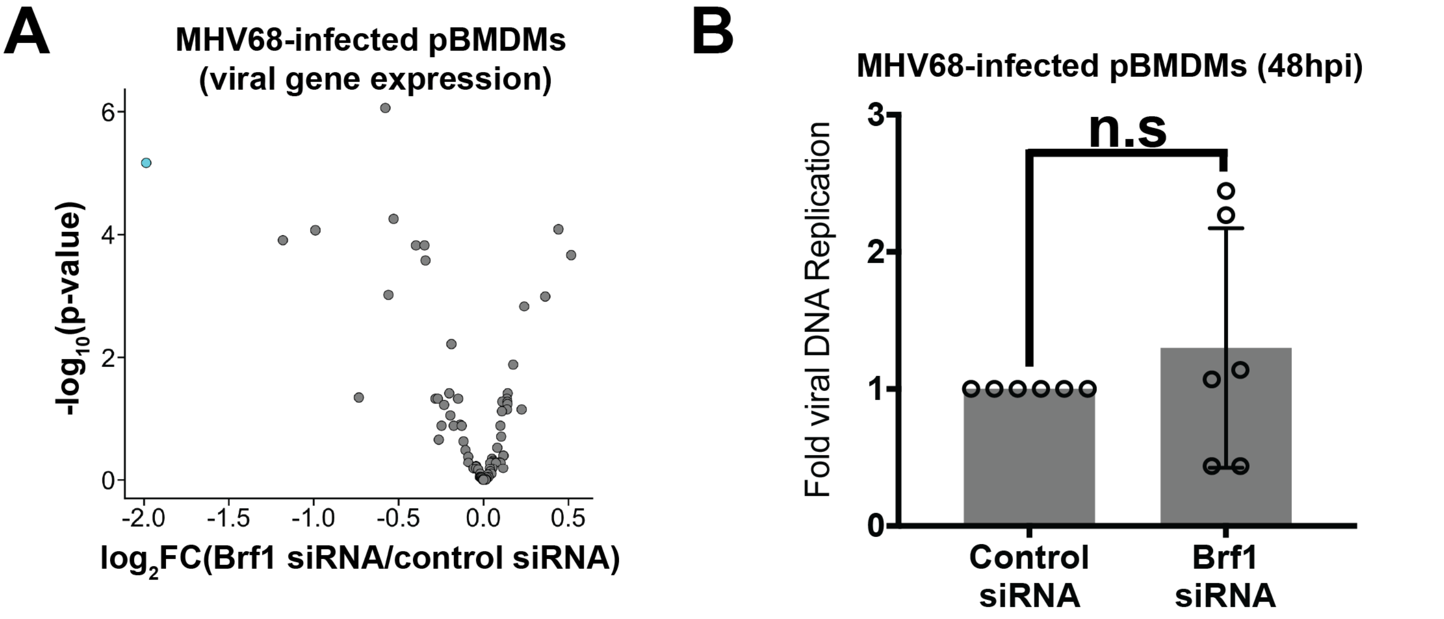


**Supplemental Figure 4. Brf1 knockdown during MHV68 infection does not impact viral gene expression or viral DNA replication in primary BMDMs. A.** Differential expression analysis at viral genes (n=79) from RNA-seq was plotted as fold change (FC) versus adjusted p-value. A p-value threshold of $<0.05$ was used to identify genes that were significantly downregulated ($log_2FC \leq -1.5$, cyan, n=1) or upregulated ($log_2FC \geq 1.5$, purple, n=0) when comparing MHV68-infected primary BMDMs transfected with Brf1 targeting siRNAs to those transfected with control non-targeting siRNA. **B.** Viral DNA replication was measured by qPCR. Primary BMDMs transfected with control non-targeting or Brf1 targeting siRNAs were infected with MHV68 at an MOI of 5 in triplicate. At 48 hours post-infection (hpi), cells were harvested and lysed to extract total DNA. Viral DNA copies were normalized to GAPDH promoter DNA and compared to values from control siRNA-transfected cells. Error bars show the standard deviation (SD), and statistics were calculated using an unpaired *t*-test on raw $\Delta C_T$ values. n.s, not significant.

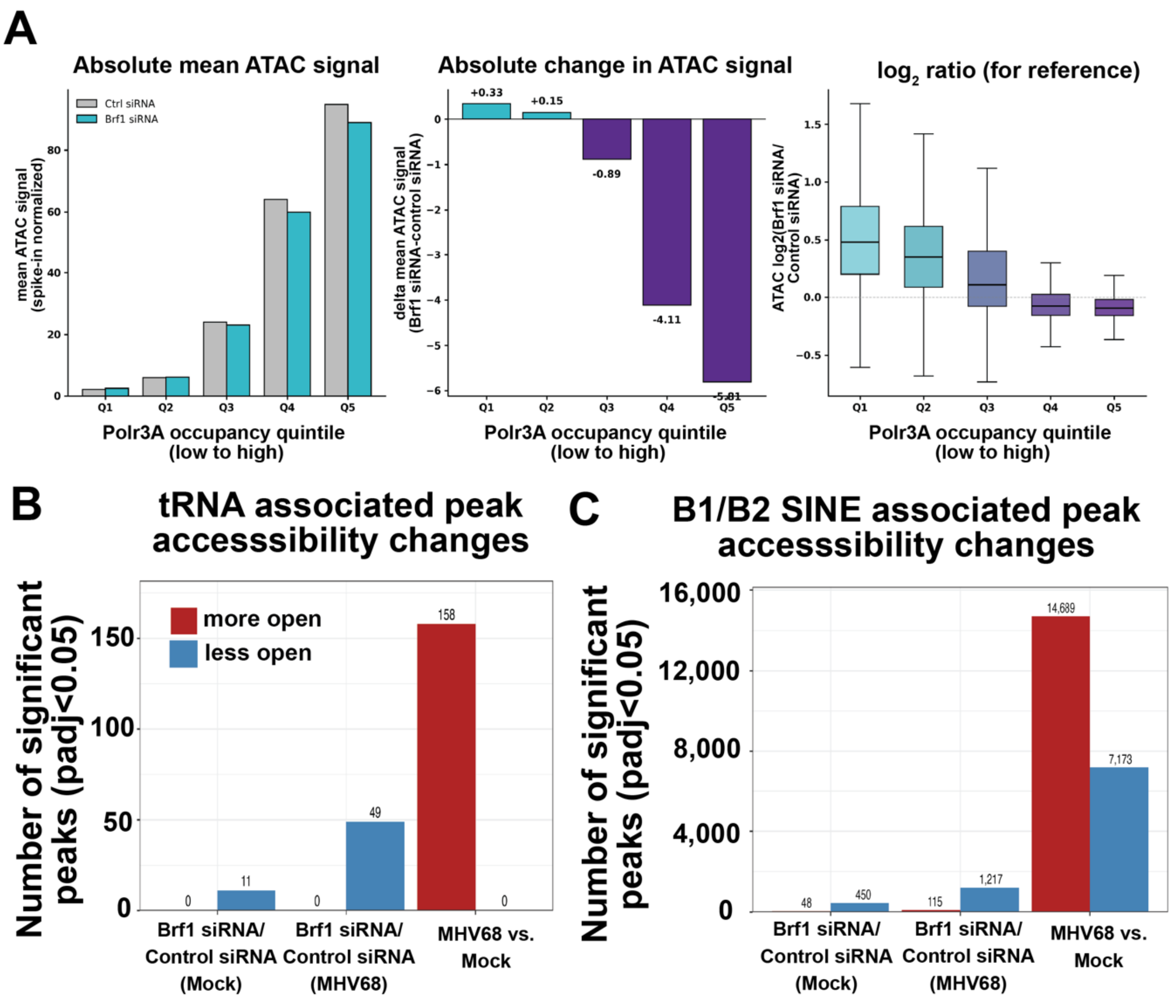


**Supplemental Figure 5. Brf1 depletion reduces chromatin accessibility at canonical Pol III targets, while MHV68 infection drives extensive opening of SINE elements. A.** Absolute mean ATAC-seq signal per Polr3A occupancy quintile in control siRNA (grey) and Brf1 siRNA (teal) NIH3T3 fibroblasts in the mock condition. Bars represent mean spike-in-normalized ATAC signal across protein-coding gene promoters (TSS ±500 bp) within each Polr3A quintile (n = ~4,000 promoters per quintile; n = 20,012 total). The top quintile (Q5; highest Polr3A occupancy) shows a reduction in absolute mean ATAC signal upon Brf1 depletion (94.9 → 89.0; −6.1%), confirming that the chromatin accessibility loss at Polr3A-occupied promoters observed in Fig.3b reflects a reduction in absolute accessibility rather than a normalization-driven artifact. The progressive dose-response across quintiles is consistent with a direct, Polr3A-occupancy-dependent role for Pol III in maintaining chromatin accessibility. **B.** Number of significantly differentially accessible tRNA-associated ATAC-seq peaks (adjusted $P < 0.05$) across three pairwise comparisons in NIH3T3 fibroblasts: Brf1 siRNA vs. control siRNA in mock conditions (left), Brf1 siRNA vs. control siRNA during MHV68 infection (middle), and MHV68 vs Mock in control siRNA cells (right). Red bars, peaks are significantly more open in the first condition; blue bars, peaks are significantly less open. Peak counts are annotated above each bar. Differential accessibility was computed using DESeq2 on a consensus ATAC-seq peak set intersected with high-confidence tRNA gene annotations (GtRNAdb v2(Chan and Lowe, 2016)). **C.** As in (B) but for ATAC-seq peaks overlapping B1 or B2 SINE retrotransposon elements (RepeatMasker B1_Mm and B2_Mm subfamilies (Smit, 2013-2015)). Brf1 depletion selectively reduces SINE accessibility in both mock and MHV68 conditions; MHV68 infection drives extensive opening of SINE elements (14,689 peaks more open), consistent with virus-induced Pol III redistribution to SINEs (Figure 1).

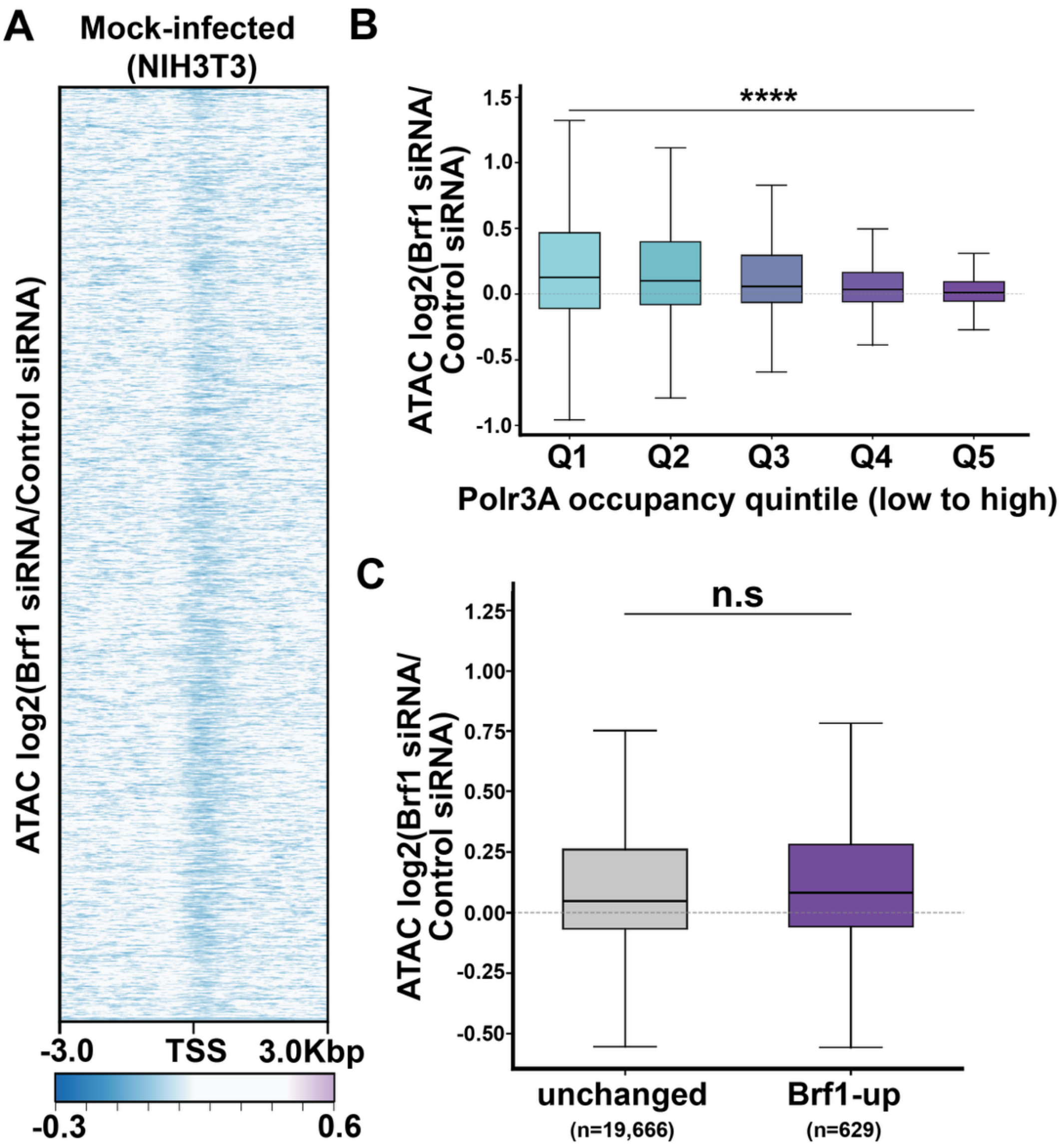


**Supplemental Figure 6. ATAC-seq analysis during MHV68 infection. A**. Heatmap of ATAC-seq $\log_2$(Brf1 siRNA/control siRNA) at protein-coding gene transcription start sites (TSSs; ±3 kb) in MHV68-infected (NIH3T3 fibroblasts; n = 20,012 protein-coding genes; n = 3 biological replicates per condition). Each row is a single protein-coding gene TSS; rows are sorted by mean $\log_2$FC. Color scale: $\log_2$(Brf1 siRNA/control siRNA); purple, accessibility gain; teal, accessibility loss; white, no change (clipped at ±0.6). Top: aggregate mean $\log_2$FC profile across all TSSs. ATAC-seq signal is from spike-in-normalized replicate-averaged BigWig files. **B.** ATAC-seq $\log_2$(Brf1 siRNA/control siRNA) at protein-coding gene promoters (TSS ±500 bp) in MHV68-infected NIH3T3 fibroblasts (48 hpi), stratified by Polr3A occupancy quintile in the MHV68 condition. Each box represents ~4,000 promoters (n = 20,295 total promoters after low-signal filtering). Boxes show median and interquartile range; whiskers show 1.5 × IQR; outliers omitted for clarity. Statistical comparison of Q5 vs Q1 distributions: two-sided Mann–Whitney U test, **** $P < 10^{-4}$. **C.** ATAC-seq $\log_2$(Brf1 siRNA/control siRNA) at promoters of Pol III–restrained inflammatory genes (n = 629 mapped TSSs from the 373 genes upregulated by Brf1 depletion during MHV68 infection; Fig. 4) compared to all unchanged genes (n = 19,666), in MHV68-infected fibroblasts. Boxes show median and interquartile range; whiskers show 1.5 × IQR; outliers omitted for clarity. Two-sided Mann–Whitney U test, n.s = not significant.

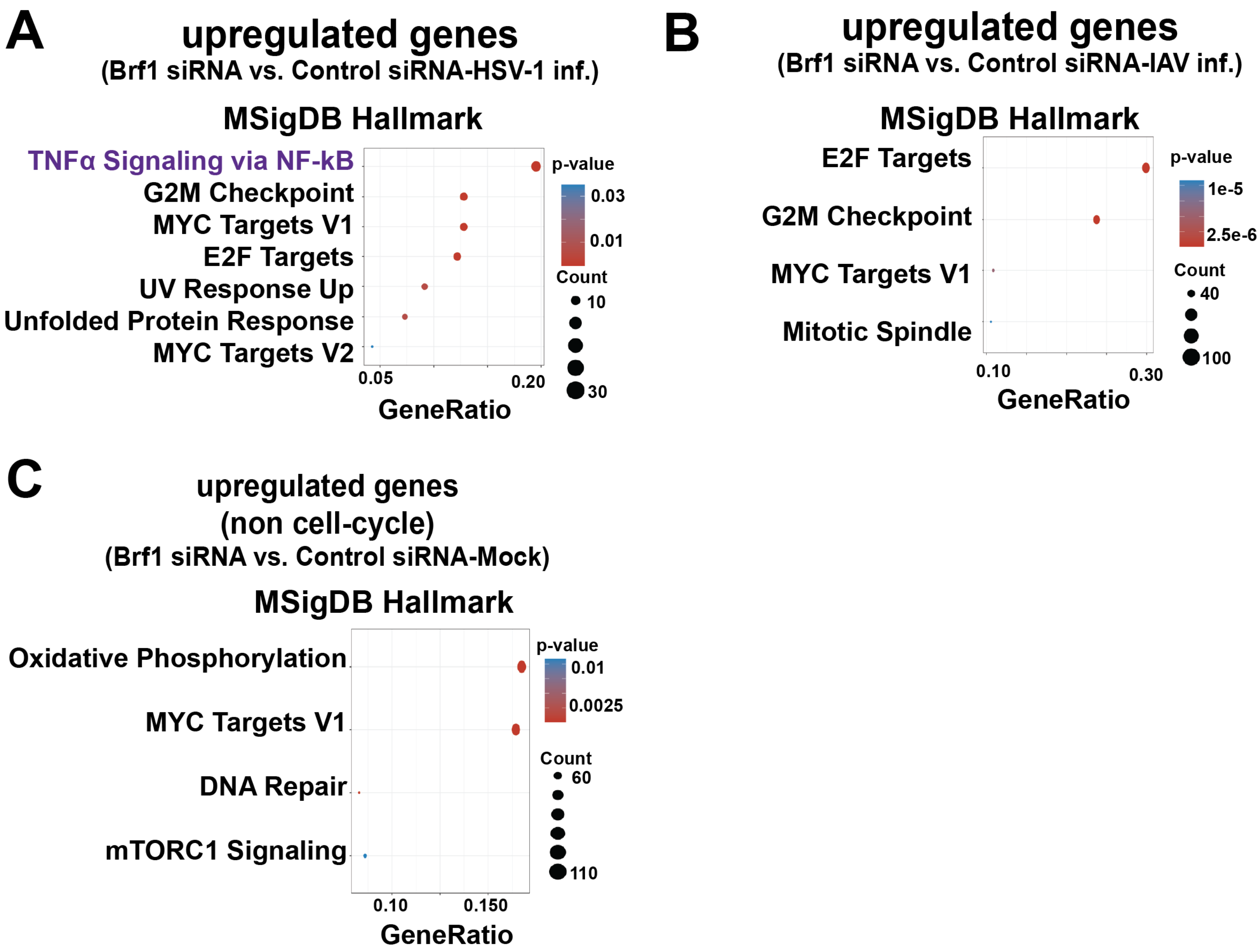


**Supplemental Figure 7. Hallmark gene set enrichment analyses of genes de-repressed by Brf1 depletion in human A549 cells. A.** MSigDB Hallmark gene-set enrichment for the full set of genes upregulated upon Brf1 depletion in HSV-1-infected A549 cells (Brf1 siRNA vs control siRNA; n=532 upregulated genes). Enrichment was performed using clusterProfiler::enricher with the human MSigDB Hallmark collection (msigdbr collection "H"); Benjamini-Hochberg multiple-testing correction; p-value cutoff 0.05. Dot position on the x-axis indicates gene ratio; dot size indicates the number of genes in each set (Count); dot color indicates adjusted p-value. **B**. As in (A), for genes upregulated upon Brf1 depletion in IAV-infected A549 cells (n = 708 upregulated genes). **C**. Hallmark gene-set enrichment for genes upregulated upon Brf1 depletion in mock-infected A549 cells, without MSigDB Hallmark E2F_TARGETS and G2M_CHECKPOINT gene-set members from both the input gene list and the tested gene-set database. This cell-cycle-excluded analysis was performed identically to that shown for HSV-1 and IAV in Figure 6C-D and serves as an infection-specificity control.
